# Structural snapshots along K48-linked ubiquitin chain formation by the HECT E3 UBR5

**DOI:** 10.1101/2023.06.06.543850

**Authors:** Laura A. Hehl, Daniel Horn-Ghetko, J. Rajan Prabu, Ronnald Vollrath, D. Tung Vu, David A. Pérez Berrocal, Monique P. C. Mulder, Gerbrand J van der Heden van Noort, Brenda A. Schulman

## Abstract

Ubiquitin chain formation by HECT catalytic domain-containing E3 ligases regulates vast biology, yet the structural mechanisms remain unknown. We employed chemistry and cryo-EM to visualize stable mimics of the intermediates along K48-linked ubiquitin chain formation by the human E3, UBR5. The structural data reveal a ≈620 kDa UBR5 dimer as the functional unit, comprising a scaffold with flexibly-tethered ubiquitin-binding UBA domains, and elaborately arranged HECT domains. Chains are forged by a UBA domain capturing an acceptor ubiquitin, with its K48 lured into the active site by numerous interactions between the acceptor ubiquitin, manifold UBR5 elements, and the donor ubiquitin. The cryo-EM reconstructions allow defining conserved HECT domain conformations catalyzing ubiquitin transfer from E2 to E3, and from E3. Our data show how a full-length E3, ubiquitins to be adjoined, E2, and intermediary products guide a feed-forward HECT domain conformational cycle establishing a highly efficient, broadly targeting, K48-linked ubiquitin chain forging machine.

## INTRODUCTION

The ubiquitin (Ub) code depends on specific types of polyubiquitin chains determining fates of modified proteins^1, 2^. For example, widespread regulation is achieved by K48-linked Ub chains triggering proteasomal degradation, while chains linked between Ub’s N-terminus or K63 are associated with signaling and membrane protein trafficking^3–7^. Branched chains, where one type of Ub chain is further modified by K48-linkages, are particularly potent at eliciting protein turnover^8–13^. Given the fundamental roles of polyubiquitin chains in biological regulation – and of K48-linked chains in particular – it is important to understand the structural mechanisms by which these specific linkages are forged.

Polyubiquitylation is achieved by E2 and E3 enzymes collaborating to link the C-terminus of a “donor” Ub (Ub^D^) to a specific site on an “acceptor” (Ub^A^). The underlying mechanism depends on the type of E3 ligase catalytic domain^14–16^. For many E3s, RING domains bind and activate Ub^D^ transfer from the catalytic Cys of an E2 to a recruited Ub^A^. As such, RING E3 linkage-specificity is typically determined by the E2 partner^17^. On the other hand, E3s with HECT and RBR catalytic domains directly mediate Ub transfer and determine the type of polyubiquitin chain produced^6, 18–20^. HECT- and RBR-family E3s have active site cysteines that generate Ub chains through two reaction steps^21–23^. A first transition state (TS1) directs Ub^D^ transfer from an E2’s catalytic Cys to that of the HECT or RBR E3. This first step produces a reactive E3∼Ub^D^ intermediate, with the C-terminus of Ub^D^ thioester-bonded to the E3. A distinct, second transition state (TS2) directs Ub^D^ transfer from the E3 Cys to the Ub^A^. Structures of RING and RBR E3 complexes, essentially chemically-trapped in action, have shown how these classes of ligases form Ub chains with various linkages^24–28^. However, Ub chain formation has not been visualized for any HECT E3.

HECT E3s were the first family of Ub ligases discovered^21, 22^. The nearly 30 human HECT E3s regulate numerous biological processes, including transcription, metabolism and membrane protein trafficking^19, 29^. They feature a C-terminal HECT (Homologous to E6AP C-terminus) catalytic domain and variable N-terminal regions^22^. Amongst other functions, the N-terminal regions are associated with regulation by autoinhibition, activation, and substrate recognition. Meanwhile, structures of isolated HECT domains from many family members showed a conserved bilobal structure; a larger N-terminal, or “N-”, lobe binds E2, and smaller C-terminal “C-lobe” harbors the catalytic Cys^30, 31^. Prior crystal structures showed a variety of arrangements between the N- and C-lobes, indicating they are tethered by a flexible interlobe linker^31^. Two frequently-observed configurations are “L” and “Inverted-T”, named based on the overall shape oriented with the long axis of the N-lobe on the bottom. These differ by an interlobe rotation of ≈150°, placing the C-lobe either to one side (L) or towards the middle of the N-lobe (Inverted-T). Although prior crystal structures have represented HECT domains at various stages of ubiquitylation cascades, no stage has been visualized more than once^32–35^. Also, there are no structures representing any Ub transfer intermediate for a full-length HECT E3, nor structures representing two different intermediates for a single E3. As such, whether or not HECT E3s mediate Ub transfer through conserved catalytic architectures remains unclear, and the functional relevance of different N-/C-lobe arrangements is the subject of debate^31, 36^.

A HECT ligase of emerging importance is the 2799-residue, multidomain, human E3 UBR5^29, 37–45^. UBR5 specifically generates K48-linked Ub chains, including by branching pre-formed K11- or K63-linked chains^9, 11^. As such, UBR5 directs degradation of diverse proteins, in part by targeting unpartnered subunits otherwise found in protein complexes^45^. UBR5 plays key roles in stem cell pluripotency, tumor suppression, oncogenesis and other important biological processes ^39–41, 45^. Prior structure-function studies showed UBR5’s UBA domain binds Ub, and suggested a role for this interaction during polyubiquitylation^46–50, 51 52^. However, without structural data showing a full-length HECT E3 in the transition states, knowledge of how the domains would be arranged across the cascade where Ub^D^ is transferred from an E2 to the HECT domain catalytic Cys, and then to a Ub^A^ remains rudimentary.

Here, we address this problem with a suite of cryo-EM reconstructions for chemically-stable proxies for the TS1, UBR5∼Ub^D^, and TS2 intermediates. Comparison to prior structures identifies a conserved HECT domain conformational trajectory for Ub^D^ transfer from E2 to the E3 to a target, while illuminating how the conformations and structural transitions along the polyubiquitylation cascade are achieved synergistically by elements of UBR5’s N-terminal regions, the HECT domain, the E2∼Ub intermediate, and the donor and acceptor Ubs. Furthermore, the structure of the TS2 intermediate shows an intricate web of interactions converging to place the acceptor Ub’s K48 into the ubiquitylation active site. Together, the data reveal a HECT E3 linkage-specific polyubiquitylation cascade, for K48 chains forged by UBR5.

## RESULTS

### Cryo-EM structure shows a dimeric UBR5 scaffold supporting HECT-mediated K48-linked Ub chain formation

HECT E3 enzymatic cascades can be examined by monitoring Ub transfer starting with an E2∼Ub intermediate (Figure 1a, ∼ refers to covalent bond to E2 or E3 catalytic Cys). A pulse reaction enzymatically generates the thioester linkage between E2 (here UBE2D2) and the donor Ub (here termed Ub^D^). Ub^D^ can be tracked by fluorescent labeling, and a K48R mutation can be employed to prevent its use as an acceptor. After the pulse reaction is quenched, E2∼Ub^D^ is added to UBR5 and unlabeled Ub^A^. Fluorescent Ub^D^ is tracked as it is transferred from E2 to E3 to Ub^A^ based on electrophoretic mobility by SDS-PAGE. This assay confirmed that wild-type (WT), full-length UBR5 expressed in HEK293S cells displayed active site Cys2768-dependent Ub chain-forming activity (Figure 1b).

**Figure 1:**
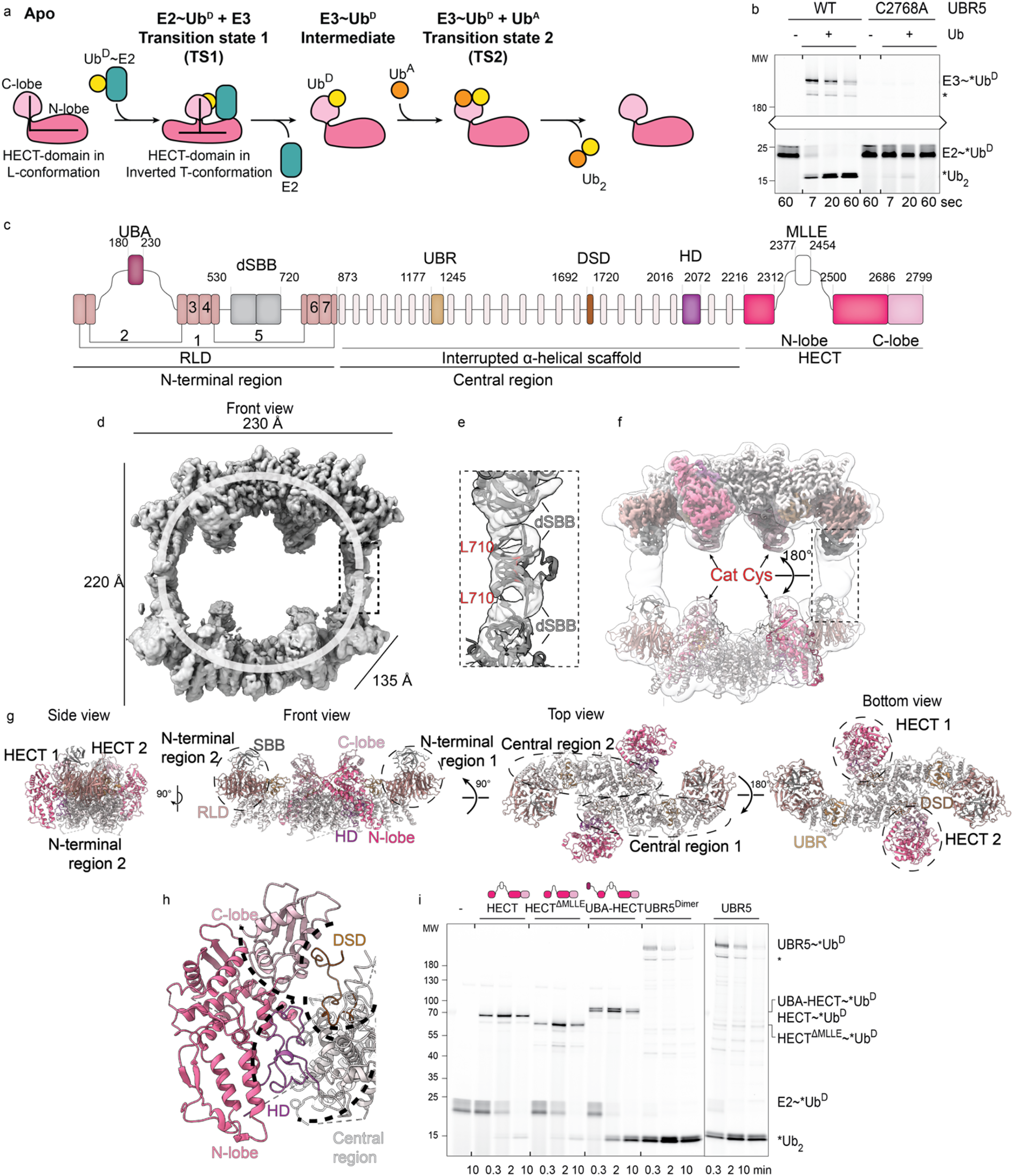
Cryo-EM of UBR5 reveals oligomeric scaffold tethering to UBA domains and elaborately arranging HECT domains. **a,** Cartoon showing HECT E3-mediated Ub chain formation step-by-step. **b,** Assay testing linkage of two Ubs into a chain by full-length WT and C2768A mutant UBR5. The assay initiates by forming the E2∼Ub^D^ intermediate, here with the E2 UBE2D2 and fluorescently-labeled K48R Ub to prevent use as acceptor (*Ub^D^). E2∼*Ub^D^ is incubated with indicated version of UBR5 and unlabeled acceptor (Ub^A^, here Ub-6xHis) for various times, and *Ub^D^ is tracked through cascade by migration in non-reducing SDS-PAGE. Hereafter, this experimental format is referred to as a di-Ub synthesis assay. A UBR5-degradation-product that remains catalytically active causes a second lane below E3∼*Ub^D^ and is marked with “*” throughout this study. **c,** UBR5 domains based on cryo-EM structure. **d,** Cryo-EM map of UBR5^C2768A^, highlighting the two U-shaped units. Dotted box indicates dSBB domains highlighted in **e** as mediating tetramerization between the two dimeric U-shaped units. **e,** AlphaFold2-model of dSBB domains showing key location of L710, in red. **f,** Transparent low-pass filtered map of tetrameric UBR5^C2768A^ is superimposed on UBR5^Dimer^ density in top half and experimentally-derived coordinates in lower half. The 180° rotation across the two halves is indicated. Dotted box corresponds to dSBB domains, not well-resolved in the UBR5^Dimer^ density. **g,** Four views of UBR5^Dimer^ structure, termed “Side”, “Top”, “Front”, “Bottom”, used to showcase different regions in subsequent figures. **h,** Close-up showing DSD and HD domain interactions with HECT domain C- and N-lobes, respectively. **i,** Di-Ub synthesis assay as in **b**, testing various versions of UBR5: structurally-redefined HECT domain with or without (HECT^ΔMLLE^) MLLE insertion, or with the UBA domain connected by a 15-residue linker, or dimeric or WT full-length UBR5.

Cryo-EM data for catalytically-inactive UBR5^C2768A^ showed a giant ≈230 Å x 220 Å x 135 Å ovoid multidomain assembly of two U-shaped units (Figure 1c-d, Extended Data Figure 1, Extended Data Table 1) 3D classification, and fitting models of domains generated by AlphaFold2 into the best of the maps, refined at 3.7 Å overall resolution, revealed four key structural properties. First, the U-shaped assembly appeared to be the fundamental structural unit, varying in angles relative to each other by up to ≈20° in different classes (Extended Data Figure 2a). Second, each U-shaped unit corresponds to a dimer of UBR5 protomers, which consist of N-terminal and central regions wherein multiple domains form a scaffold supporting the C-terminal HECT domain (Figure 1c, Extended Data Figure 2b). The N-terminal region consists of an interrupted RLD β-propeller, with UBA (not visible in the map) and dSBB (double ***<u>S</u>***mall ***<u>B</u>***eta ***<u>B</u>***arrel) domains inserted in blades 2 and 5, respectively (Extended Data Figure 2c). The central region is primarily ⍺-helical, but also embeds the potential substrate-binding UBR domain (Extended Data Figure 2d) and other elements. The two HECT domains project towards the center of each U-shaped dimer from opposite directions. Finally, within the tetrameric oval, there are two dimerization interfaces. Two protomers are rigidly affixed by extensive interactions across their central regions at the center of the U (Extended Data Figure 2e). Meanwhile, two dimers are more loosely connected in the oval, by interactions between dSBB domains at the tops of the “U”s. Structural modeling suggested an L710D mutation at the dSBB interface could disrupt the tetrameric assembly and allow production of a stable dimer (Figure 1e). Indeed, mass photometry confirmed WT UBR5 preferentially forms a tetramer, and the L710D mutant (hereafter referred to as UBR5^Dimer^) primarily forms a dimer (Extended Data Figure 2f).

Cryo-EM data obtained for UBR5^Dimer^ yielded a map allowing building and refining a 2-fold symmetric experimentally-derived structure at 2.7 Å resolution (Figure 1f-g, Extended Data Figure 2g-i, Extended Data Figure 3, Extended Data Table 1). The structures and speculated functions of the individual domains comprising the UBR5 scaffold are extensively discussed in manuscripts that were posted on bioRxiv during the preparation of ours^50, 52^. UBR5’s C-terminal HECT domain adopts the L-conformation (Extended Data Figure 2h). As in most crystal structures of isolated HECT domains, UBR5’s six C-terminal “tail” residues are not visible and are presumably disordered^31^. Here, we focus on the elements establishing the architecture of the HECT domain, and the structural basis of polyubiquitylation.

UBR5 displays several unique HECT domain features. The HECT domain N-lobe is interrupted by insertion of the MLLE domain^47, 53^, which has been implicated in binding to some substrates but is not visible in any of our cryo-EM maps. As such, part of the HECT domain was excluded from prior annotations^54^. The structure redefines UBR5’s HECT domain as corresponding to residues 2216-2312 and 2500-2799 (Extended Data Figure 2h).

Second, the HECT domain L-configuration is stabilized by multiple elements emanating from the central region of the scaffold. The central region both mediates dimerization, and contains two meandering sequences – which we term DSD (Domain Swap Dimerization) and HD (HECT Display) domains – that bind the HECT domain. As part of the extensive dimerization interface within the scaffold, one protomer’s DSD domain is partly embedded in a groove of the other subunit. A peptide-like loop from the DSD domain extends beyond the scaffold to bind the C-lobe from one HECT domain (Extended Data Figure 2i). Meanwhile, one side of the 6 kDa HD domain interacts with the ⍺-helical portion of the scaffold, and the other side nestles in a concave surface of the N-lobe. Because the residues leading to and from the HD domain are not visible in the EM density, it could in principle rotate to display the HECT domain N-lobe in various orientations (Extended Data Figure 2j).

The specific arrangement of UBR5’s HECT domain is established through seven-way interactions: between the C-terminus of the central region and N-terminus of the HECT domain, between the DSD domain and HECT domain C-lobe, between the central region and HD domain, between the HD domain and HECT domain N-lobe surface opposite the canonical E2-binding site, between the HECT domain N- and C-lobes, between the ⍺-helical portion of the scaffold and the DSD domain, and between the DSD and HD domains. The elaborate nature of the assembly portended an important role of this specific HECT domain architecture in UBR5-mediated polyubiquitylation (Figure 1h).

Given that many isolated HECT domains are sufficient to mediate Ub chain formation^19, 30, 31^, we assayed the structurally-redefined HECT domain side-by-side with FL UBR5 and UBR5^Dimer^. Much like WT UBR5, UBR5^Dimer^ retained robust Ub chain forming activity (Extended Data Figure 2k), and specificity for forging K48 linkages (Extended Data Figure 2l). However, truncated versions corresponding to the structurally-redefined UBR5 HECT domain, with or without the MLLE domain insertion, produced little di-Ub (Figure 1i). We considered that the UBA domain could play a key role, because other Ub chain forming enzymes depend on Ub-binding domains to recruit an acceptor Ub^24–28, 55^. A minimal version of UBR5 with the UBA and HECT domains connected by a 15-residue linker did generate di-Ub chains, but less efficiently than UBR5 and UBR5^Dimer^. Taken together, the data showed the UBR5 UBA and HECT domains play critical roles in Ub chain formation, and that its high level of activity makes UBR5^Dimer^ suitable for structurally defining the Ub chain-forming cascade.

### Visualizing a stable mimic of the transition state during Ub transfer from E2 to UBR5

We sought structural insights into the TS1 catalytic assembly, which mediates transfer of Ub^D^ from the E2’s active site Cys to the E3’s active site Cys (Figure 2a). A stable mimic of this otherwise fleeting reaction was generated with the E2 UBE2D, by adapting our method that allowed visualizing the TS1 intermediate for an RBR E3^56^. An electrophile installed at the active site of a proxy for the UBE2D∼Ub intermediate reacted with UBR5^Dimer^, dependent on the catalytic Cys (Extended Data Figure 4a-c). Cryo-EM of the UBR5^Dimer^∼Ub^D^∼UBE2D complex showed considerable heterogeneity (Extended Data Figure 5). Nonetheless, a map refined without symmetry at 7.3 Å overall resolution showed density for one HECT∼Ub^D^∼E2 domain assembly (Figure 2b). A structural model was generated by fitting the map with prior coordinates for UBE2D∼Ub^32^, and from three units from the UBR5^Dimer^ cryo-EM structure: the scaffold; the HD domain and HECT domain N-lobe; and HECT domain C-lobe (Figure 2c).

**Figure 2:**
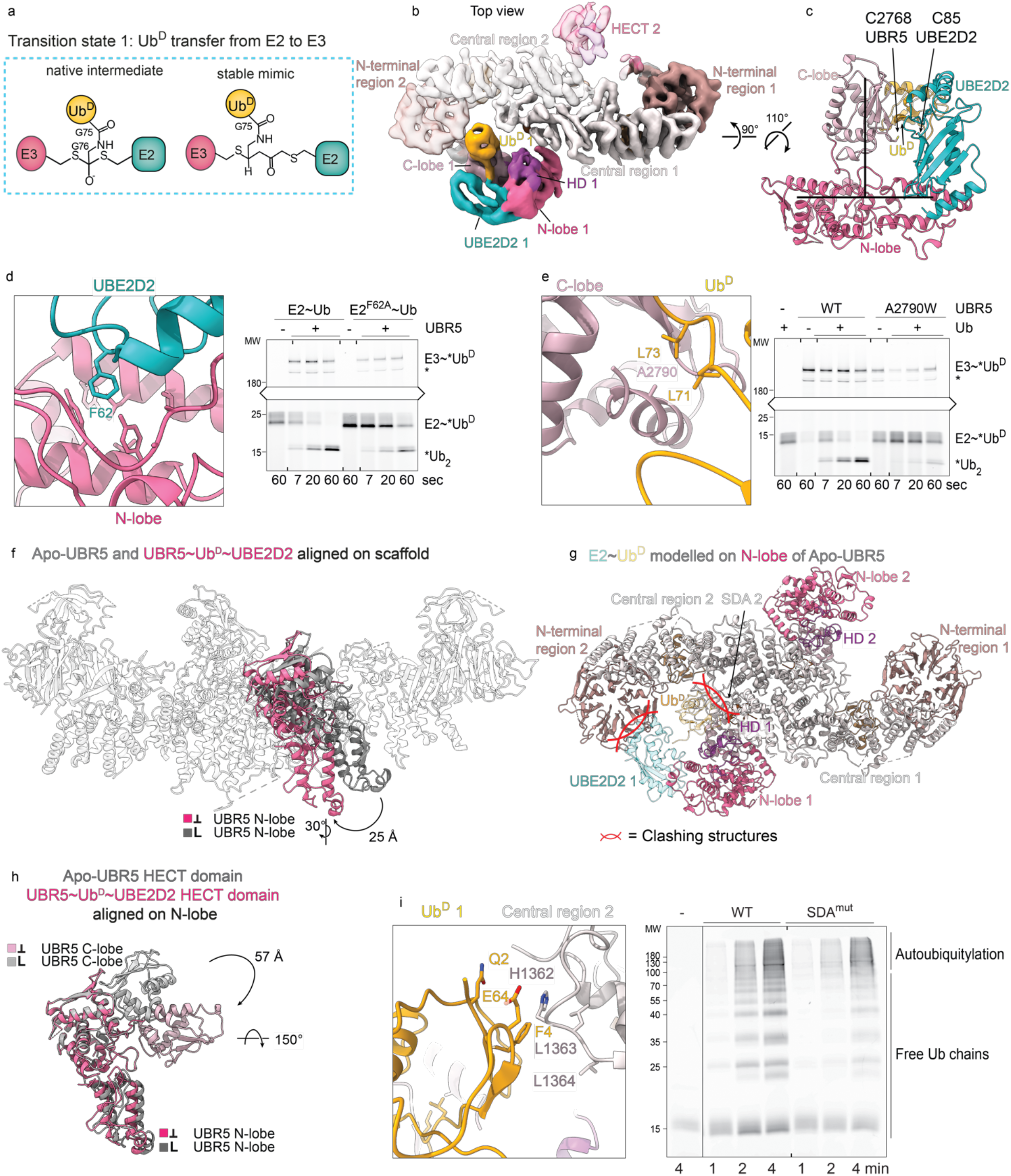
Cryo-EM map visualizing Ub^D^ transfer from E2 to UBR5. **a,** Chemical structures of native transition state 1 (TS1) for Ub transfer from the E2 UBE2D2 to UBR5, and chemically-stable mimic. The TS1 intermediate is mimicked by installation of an electrophilic moiety between the C-terminus of Ub (truncated at G75, representing Ub^D^) and UBE2D2’s catalytic Cys, which traps UBR5^Dimer^’s catalytic Cys via a stable 3-way crosslink. **b,** Cryo-EM map of UBR5^Dimer^∼Ub^D^∼UBE2D2 in “top” view shown in Figure 1g. **c,** Close-up of UBE2D2∼Ub-bound UBR5 HECT domain. The structural model was made by by individually fitting the N-lobe and C-lobe of UBR5^Dimer^, and published coordinates for UBE2D2∼Ub (PDB: 3JWZ) into the density. The HECT domain was modelled by individually fitting the N-lobe and C-lobe of UBR5^Dimer^. Locations of E2 (UBE2D2 C85) and E3 (UBR5 C2768) active site cysteines are labeled. **d,** Left, close-up of UBR5^Dimer^∼Ub^D^∼UBE2D2 structural model over E2-N-lobe interface, highlighting UBE2D2’s F62 docking in UBR5s aromatic pocket. Right, di-Ub synthesis assay testing role of interface by comparing WT or F62A mutant E2 (UBE2D3 in assay). **e,** Left, close-up of UBR5^Dimer^∼Ub^D^∼UBE2D2 structural model over HECT domain C-lobe interface with Ub^D^, highlighting UBR5 A2790 inserted in Ub^D^ hydrophobic pocket comprising L71 and L73. Right, di-Ub synthesis assay testing effect of A2790W mutant designed to disrupt interface. **f,** Structural superposition of apo or E2∼Ub^D^-engaged UBR5^Dimer^ structures aligned on the scaffold show their distinctly positioned HECT domain N-lobes in grey and pink, respectively. The C-lobes and E2∼Ub^D^ are not shown. The colored squares indicate the L- and inverted-T-conformations of the HECT domain in the two complexes, and the respective coloring. This depiction of the respective conformation and coloring will be used throughout the study. **g,** Requirement for reorientation of the HECT domain N-lobe relative to the scaffold to achieve the TS1 reaction is shown by docking E2∼Ub^D^ on N-lobe of apo UBR5. E2 and Ub cannot bind the apo-UBR5 conformation because these would clash with the N-terminal and central regions of the scaffold. Clashing regions are indicated with intersecting red arcs. **h,** Relative rotation of the C-lobe between the apo UBR5 (grey) and TS1 (pink) conformation is shown by aligning the HECT domain on the N-lobe. L- and Inverted T-conformation of the respective complex as well as the color-coding are indicated with colored boxes and symbols. **i,** Left, close-up of UBR5^Dimer^∼Ub^D^∼UBE2D2 structural model showing UBR5 Scaffold-Donor Ub-Approaching (SDA) loop (residues H1362-L1364) and the F4-patch of Ub^D^. Right, polyubiquitylation assay testing effect of SDA-mutant (H1362D L1363D L1364D). Assay was performed by mixing E1, E2 (UBE2D2), UBR5, and fluorescent Ub (*Ub) for indicated times prior to resolving ubiquitylated species on reducing SDS-PAGE gels.

The model shows Ub^D^’s C-terminus and the active site cysteines of UBE2D and UBR5^Dimer^ juxtaposed as expected for the TS1 intermediate. Notably, the HECT domain N- and C-lobes adopt the Inverted-T, not the L, conformation, placing the E2 and E3 active sites for Ub transfer between them. The T-conformation is stabilized by avid interactions between the HECT domain and the E2∼Ub^D^ intermediate. The E2 binds the HECT domain N-lobe, while its catalytic Cys-linked Ub engages the HECT domain C-lobe. Accordingly, mutating a key UBE2D residue binding UBR5’s N-lobe (F62A)^57^, or the key UBR5 C-lobe residue at the noncovalent interface with Ub^D^ (A2790W), impaired E3 ligase activity (Figure 2d-e).

While the scaffold itself superimposes between the apo and E2∼Ub^D^-bound UBR5 assemblies, there are substantial differences in the relative orientations of the HECT domain N- and C-lobes. First, the N-lobe tilts by 25° relative to the scaffold (Figure 2f). This rotation is required because the N-lobe position in apo UBR5 is incompatible with E2∼Ub^D^ binding: a bound E2 would clash with the RLD from the opposite protomer in the dimer, and its linked Ub^D^ would clash with the ⍺-helical domain from one protomer and a loop we term “SDA” (Scaffold Donor-Ub Approaching) from the other (Figure 2g). At this resolution, we cannot unambiguously determine if UBR5’s N-lobe remains bound to the HD domain in the TS1 intermediate, although the necessary rotation could be achieved by release of some interactions with the scaffold (Extended Data Figure 4d). Second, in order to face the N-lobe-bound E2, the C-lobe has rotated ≈150° about the interlobe tether (Figure 2h). This shifts the position of the E3 catalytic Cys by >40 Å. This positioning of the C-lobe grants UBR5’s catalytic Cys access to the E2∼Ub^D^ active site, but would require its disengagement from the DSD domain. Thus, it seems that E2∼Ub^D^ binding not only directs the catalytic architecture of the HECT domain, but also orchestrates substantial rearrangement in the context of the UBR5 scaffold.

Finally, in the UBR5∼Ub^D^∼E2 assembly, the donor Ub F4 patch is poised to graze the SDA-loop (residues H1362-L1364). We tested the function of the SDA loop by replacing the sequence with aspartates. The SDA mutation did not overtly affect Ub^D^ transfer from E2 to UBR5, nor to an acceptor Ub in pulse-chase assays monitoring di-Ub synthesis (Extended Data Figure 4e). However, the SDA loop mutant showed a subtle but obvious defect in forming low-molecular weight polyUb chains in assays with multiple cycles of E1-E2-UBR5 activities (Figure 2i).

### Cryo-EM analyses of a chemically-stable mimic of the UBR5∼Ub^D^ intermediate

After its transfer from the E2, Ub^D^ is thioester-bonded to UBR5’s catalytic Cys (Figure 3a). We generated a stable proxy for this intermediate by mixing Ub-vinyl methyl ester (Ub-VME)^58^ with UBR5^Dimer^, which reacted depending on the catalytic cysteine (Figure 3a, Extended Data Figure 4f-g). Cryo-EM of the resultant UBR5^Dimer^∼Ub^D^ complex yielded a map at 5.3 Å resolution (Figure 3b-c, Extended Data Figure 6, Extended Data Table 1). The map superimposed with the structure of apo-UBR5^Dimer^, with one striking difference: density for Ub^D^ adjacent to both C-lobes. We generated a structural model by wholesale docking of: (1) Ub^D^ ^33^, (2) the UBR5^Dimer^ structure through the HECT domain N-lobe, and (3) the C-lobe and DSD domain, which are slightly rotated compared to the structure of UBR5 without a covalently-linked Ub^D^.

**Figure 3:**
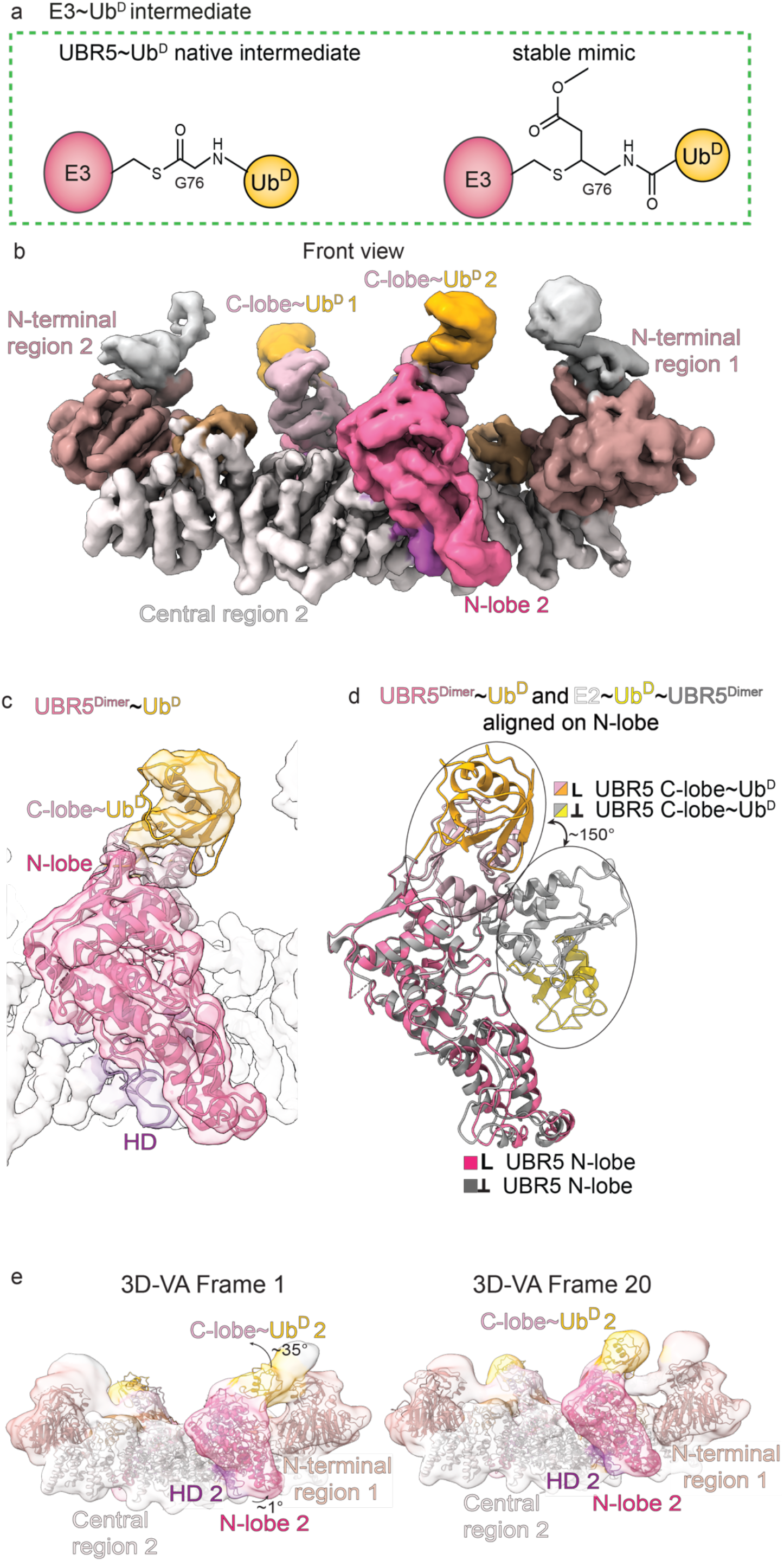
Cryo-EM of stable mimic of UBR5^Dimer^∼Ub^D^ intermediate. **a,** Chemical structures of native intermediate and chemically-stable mimic. The native TS1 reaction product is Ub^D^ thioester-bonded to UBR5’s catalytic Cys. The stable mimic used Ub-vinylmethylester (UbVME) to couple Ub^D^ to UBR5^Dimer^’s catalytic Cys. **b,** Cryo-EM map of stable mimic representing UBR5^Dimer^∼Ub^D^ in the front view. **c,** Structural model of UBR5^Dimer^∼Ub^D^ fitted into the density. **d,** Conformational change after Ub^D^ transfer from E2 to UBR5 shown in overlay of HECT domain from UBR5^Dimer^∼Ub^D^ (L conformation) in pink/orange and from UBR5^Dimer^∼Ub^D^∼UBE2D2 (Inverted-T conformation, UBE2D2 not shown) in grey/yellow. **e,** Left, frame 1 and right, frame 20 of 3D-VA performed on UBR5∼Ub^D^. The central and N-terminal region, N-lobe, C-lobe, and Ub^D^ were individually fitted into the density.

Comparing the cryo-EM-based models for UBR5^Dimer^∼Ub^D^∼E2 and UBR5^Dimer^∼Ub^D^ showed common interactions between Ub^D^ and the HECT domain C-lobe. However, in the UBR5^Dimer^∼Ub^D^ complex, the scaffold and HECT domain reverted to a configuration like apo-UBR5 (Figure 3d). The structures suggest that after Ub^D^ would be transferred from UBE2D2 to UBR5 with the HECT domain in the Inverted-T-configuration, the C-lobe and its linked Ub^D^ turn around and face the opposite direction, in the L-orientation. Such a structural progression would rely on inherent flexibility of the UBR5^Dimer^∼Ub^D^ complex. To gain insights into this structural heterogeneity, we applied 3-dimensional variability analysis (3D-VA) in CryoSPARC^59^. Examining the output conformations revealed a spectrum of orientations for the Ub^D^-linked C-lobe (Figure 3e). One extreme is intermediary between the HECT domain Inverted-T- and L-conformations, and the other extreme is the final L-orientation. The 3D-VA is consistent with a conformational progression whereby the C-lobe and its covalently-linked Ub^D^ rotate as a unit about the linker to the N-lobe.

Meanwhile, examining the 3D-VA output conformations for the overall assembly showed the scaffold and HECT domain N-lobe generally oriented as in the apo UBR5^Dimer^ structure and the model of UBR5^Dimer^∼Ub^D^. We speculate that N-lobe reorientation would be enabled by E2 dissociation after Ub^D^’s C-terminal linkage is transferred to UBR5. Elimination of constraints from E2 clashing with UBR5’s N-terminal region presumably facilitates relocation of the N-lobe, which we speculate then facilitates redirection of the Ub-bound C-lobe into the L-configuration.

### Cryo-EM structure showing mechanism of K48-linked Ub chain formation

K48-linked chains are formed through transfer of Ub^D^’s C-terminus from UBR5’s catalytic Cys to K48 on Ub^A^ (Figure 1a). To visualize the catalytic assembly, we adapted a method previously used to study deubiquitylating enzymes and RBR E3s^14, 56^. A stable mimic of the fleeting transition state (TS2) was generated as follows (Figure 4a). An electrophile was installed between the C-terminus of a truncated Ub^D^ and a Cys replacement for K48 on Ub^A^ (Extended Data Figure 7a-b). After reaction with UBR5^Dimer^, the UBR5^Dimer^∼Ub^D^∼Ub^A^ complex was purified and subjected to cryo-EM (Extended Data Figure 8, Extended Data Table 1). A 4 Å resolution cryo-EM map largely superimposed with the structure of the apo UBR5^Dimer^∼Ub^D^ complex, with minor variation in the orientation of the HECT domain C-lobe (Figure 4b). The HECT domain C-lobe binds the globular domain of Ub^D^, as observed in the cryo-EM maps for the UBR5^Dimer^∼Ub^D^∼E2 and UBR5^Dimer^∼Ub^D^ complexes. Strikingly, clear density corresponding to a second Ub could be observed next to Ub^D^ and the C-lobe of UBR5.

**Figure 4:**
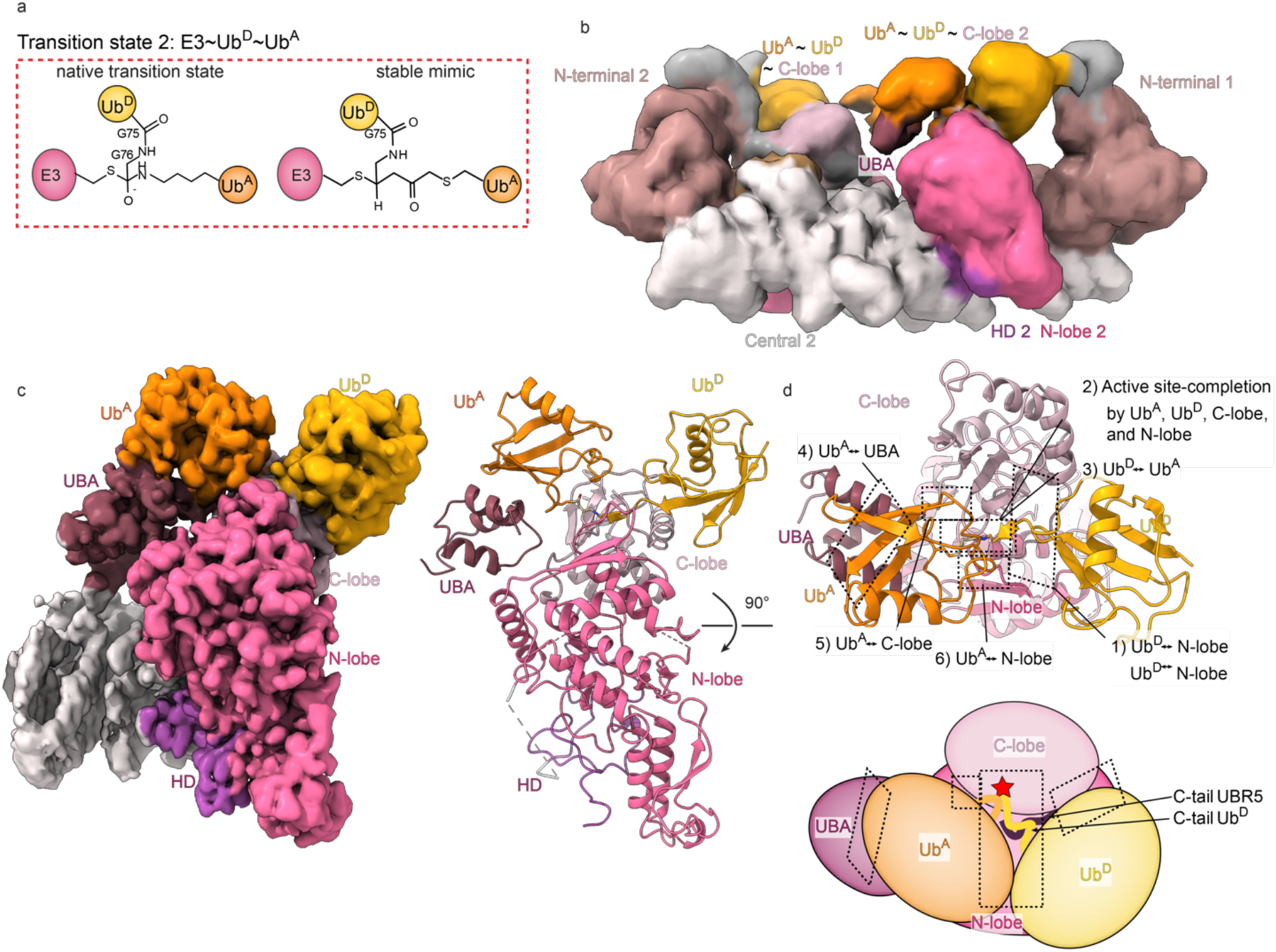
Cryo-EM structure visualizing HECT E3-mediated linkage-specific Ub chain formation. **a,** Chemical structures of native transition state 2 (TS2) for Ub chain formation, and chemically-stable mimic. The TS2 is mimicked by installation of an electrophilic moiety between the C-terminus of Ub (truncated at G75, representing Ub^D^, yellow) and a Cys replacement for the acceptor Ub’s K48 (Ub^A^, orange), which traps UBR5^Dimer^’s catalytic Cys via a stable 3-way crosslink **b,** Initial low-resolution cryo-EM map of stable mimic of the UBR5^Dimer^∼Ub^D^∼Ub^A^ complex (TS2). **c,** Local refined map and atomic model for the catalytic complex mediating K48-linked Ub chain formation, wherein the UBA domain recruits Ub^A^ and residue 48 is placed at the HECT∼Ub^D^ active site. N- and C-lobes, Ub^A^, Ub^D^, and the UBA domain are labeled. **d,** Structure above, and cartoon below, indicating interfaces establishing the catalytic geometry for K48-linked Ub chain formation.

Local refinement yielded a 3.3 Å resolution map resolving elements defining the polyubiquitylation-active state (Figure 4c-d). The interactions involving the HD and DSD domains stabilizing the HECT domain L-conformation are maintained as in UBR5^Dimer^ alone. In addition to Ub^D^ and Ub^A^, the map also shows the UBA domain and additional density around the active site that was not visible in the other maps. The UBA domain binds Ub^A^, positioned with its residue 48 at the active site. With the HECT domain in the L-conformation, the Ub^D^-linked catalytic Cys is situated not only adjacent to the acceptor, but also at the junction with the N-lobe, which is thus also poised to contribute to Ub chain formation.

Numerous previously unobserved interactions converge to form the linkage-specific Ub chain forming machinery (Figure 4d, Figure 5a). First, interactions between the HECT domain C-lobe and Ub^D^ both position the donor Ub and shape the active site (Figure 5b). The map revealed the details of the noncovalent interface between UBR5’s C-lobe and Ub^D^, observed across the cryo-EM maps for intermediates along the cascade. An intermolecular hydrophobic core is formed between UBR5’s A2790, F2732, L2762, and L2789 and Ub^D^’s Ile36, P37, L71 and L73. The hydrophobic interactions are buttressed through numerous polar interactions, involving D39, Q40, T9 of Ub^D^ and UBR5’s H2761, T2764, and K2792.

**Figure 5:**
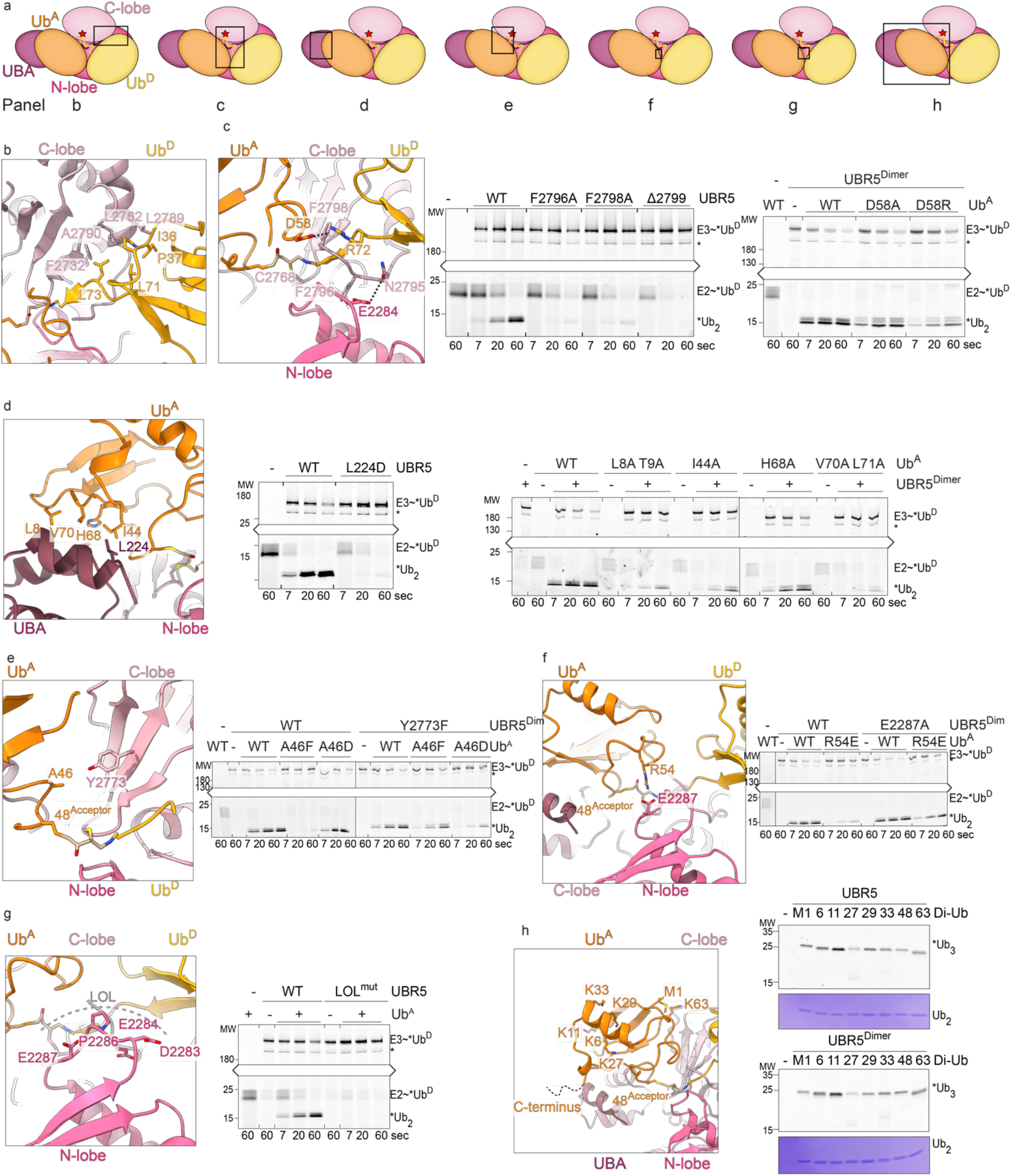
Elaborate web of interactions configuring catalytic conformation for HECT E3-mediated K48-linked Ub chain formation. **a,** Cartoon of polyubiquitylation active site, indicating regions shown on panels b-h below. **b,** Zoom-in over non-covalent interface between UBR5 C-lobe and its linked Ub^D^, also observed in lower resolution maps of TS1 and UBR5^Dimer^∼Ub^D^ intermediates. **c,** Left, close-up of multilayered active site assembly, highlighting UBR5’s C-tail residues (light pink), Ub^D^ (yellow) with its C-terminus linked to C2768 in UBR5’s C-lobe (light pink), UBR5’s N-lobe (magenta), and Ub^A^ (orange). Hydrogen bonds as well as a salt bridge are indicated with a dotted line. Right, di-Ub synthesis assays testing effects of mutating UBR5 C-tail F2796 that binds the N-lobe, penultimate F2798 that acts as a molecular glue at interface with Ub^A^, and Ub^A^ D58 that contacts Ub^D^ (Note: right-most panel uses UBR5^Dimer^ as E3). Only upper and lower portions of gels corresponding to molecular weights of *Ub^D^-linked moeties are shown, connected, for clarity. **d,** Left, close-up of UBA domain interactions with Ub^A^. Right, di-Ub synthesis assays testing effects of mutating UBR5 UBA domain and Ub^A^ interface residues. (Note: right-most panel uses UBR5^Dimer^ as E3). **e,** Left, close-up of Ub^A^ interactions with UBR5 C-lobe. Right, di-Ub synthesis assays testing effects of mutating Ub^A^ A46 at interface with C-lobe, or the opposing UBR5 residue Y2773, with mutant combinations designed to improve or impair structural compatibility. UBR5^Dimer^ was used as E3. **f,** Left, close-up of Ub^A^ interactions with UBR5 N-lobe. Right, di-Ub synthesis assay testing effects of mutating Ub^A^ R54 at interface, or UBR5^Dimer^ N-lobe mutant designed to ameliorate charge-repulsion. UBR5^Dimer^ was used as E3. **g,** Left, close-up over the ligation-organizing-loop (LOL). Right, di-Ub synthesis assay testing effecs of mutating entire LOL (residues 2283-2287) to alanines. **h,** Left, potential for Ub^A^ to be part of a Ub chain evaluated in close-up showing its C-terminus, lysines, and N-terminus that could be involved in linkages to other Ubs. Right, fluorescent scanned gels from tri-Ub synthesis assays with either WT UBR5 or UBR5^Dimer^, testing di-Ubs with the indicated linkages for capacity to serve as acceptors for *Ub^D^. Coommasie-stained gels of input di-Ubs are shown below.

Second, the active site is configured by extensive additional contacts between the HECT domain C-lobe and Ub^D^. Elements from Ub’s C-terminal tail, UBR5’s C-terminal tail, and the HECT domain catalytic loop wrap around each other as if in a 4-layered sandwich (Figure 5c). At one edge, the so-called -4 Phe^60^ (UBR5 F2796) packs between the N-lobe and Ub’s C-terminus linked to the catalytic Cys. On the other side of UBR5’s C-terminal tail, N2795 forms a hydrogen bond stabilizing the β-sheet between Ub’s C-terminus and the HECT domain C-lobe strand that culiminates at the catalytic Cys. Meanwhile, Ub^D^’s R72 and C-terminus wrap around UBR5’s penultimate F2798. F2798 in turn secures UBR5^Dimer^∼Ub^D^ active site and inserts into the interface as a molecular glue affixing Ub^A^. Accordingly, Ala substitutions for either F2796 or F2798, or projection of a negative charge into this interface through deletion of UBR5’s subsequent C-terminal residue (not visible in the map) impairs Ub chain formation.

Third, the Ub^A^ location is also positioned by an interaction with the HECT domain-linked Ub^D^ (Figure 5c). The Ub^D^ R72 – itself oriented by UBR5’s penultimate F2798 – contacts D58 from Ub^A^. Although it was not possible to test effects of R72 mutations due to requirements for this residue in generating the E2∼Ub^D^ intermediate^61, 62^, a D58R mutant Ub^A^ was defective at forming a Ub chain.

Fourth, UBR5’s UBA domain binds and presents the acceptor Ub to the active site (Figure 5d). A prior crystal structure of the isolated UBR5 UBA domain bound to free Ub was readily docked into the map^46^. This allowed visualizing details of the interaction despite relatively lower resolution density for this region. The weaker density presumably reflects conformational heterogeneity, possibly arising from the 101- and 126-residue flexible tethers between the scaffold and UBA domain that are not visible. Nonetheless, the model shows the canonical hydrophobic interactions between a UBA domain and the acceptor Ub’s I44-centered hydrophobic patch. Importantly, a UBR5 UBA domain L224D mutant, at the center of the interface, causes accumulation of the UBR5∼Ub^D^ complex, and a severe defect in Ub^D^ transfer to Ub^A^. Mutations of Ub^A^ residues contacting the UBA domain cause similar defects.

Fifth, the acceptor Ub’s residue 48 (normally K48 but here a chemically-modified Cys) is secured in the active site through interactions between adjacent residues and the HECT domain C- and N-lobes. On one side, Ub^A^’s A46 nestles opposite of Y2773 from UBR5’s C-lobe (Figure 5e). Accordingly, mutating Ub^A^’s A46 to Asp or Phe, which would hinder the structurally-observed interface, impairs di-Ub synthesis. The effects on ubiquitylation activity scale with predicted effects on the interaction. Specifically, a Ub^A^ A46F would be too large for the pocket, while A46D could potentially retain a suboptimal contact with UBR5 Y2773. On the other hand, a Y2773F mutation, which could accommodate Ub^A^’s A46, shows WT Ub chain-forming activity. The ultimate test of the importance of an interface is restoration by compensatory changes. Thus, we assayed mutant combinations predicted to improve structural compatibility. Indeed, the smaller UBR5 Y2773F side-chain partially restores activity with the Ub^A^ A46F mutant, while loss of the hydroxyl on the E3 side accounts for the further reduced activity with Ub^A^ A46D.

On the other side of Ub^A^ residue 48, its R54 projects toward E2287 in the HECT domain N-lobe (Figure 5f). Introducing a charge-repulsive R54E Ub^A^ mutation severely impairs di-Ub synthesis. We next sought a compensatory UBR5 mutant. Although we have been unable to obtain UBR5 mutant with E2287 replaced by a basic residue for technical reasons, we could test effects of eliminating the charge repulsion with an Ala mutation, which indeed partially restored di-Ub synthesis.

Finally, the HECT domain N-lobe loop comprising residues D2283-E2287, which we term the **L**igation-**O**rganizing-**L**oop (LOL), also secures the catalytic architecture by serving as a platform aligning the catalytic Cys linked to Ub^D^’s C-terminus and UBR5’s C-terminal tail (Figure 5g). Replacing the LOL sequence with alanines specifically impaired Ub chain formation. Because this loop contains three acidic residues, we also performed assays at high pH to test the structural role by offsetting potential effects on acceptor Lys deprotonation (Extended Data Figure 7c). The loop mutant was defective in all conditions tested, consistent with a role in structurally organizing the ubiquitylation active site.

### Catalytic architecture accommodates branched Ub chain formation

UBR5 has been implicated in generating branched polyUb chains in cells^9, 11^. Branched Ub chain formation could occur through UBR5 transferring Ub^D^ to a Ub^A^ within another chain. The UBR5^Dimer^∼Ub^D^∼Ub^A^ structure shows how K48 could be presented from various chain types within the transition state (Figure 5h). Ub^A^’s C-terminal tail – not visible in the map – points away from the catalytic assembly. This shows that the distal Ub in diverse chains – including K48-linked chains – would be readily modified by UBR5. Amongst Ub^A^’s lysines and the N-terminus, only residue 48 is fully buried by the catalytic assembly. Nonetheless, K11 stands out as most distal from the active site, and with its primary amino group fully exposed. This is notable because UBR5 was implicated in production of a major fraction of cellular K11/K48 branched Ub chains^9^. Consistent with the UBR5^Dimer^∼Ub^D^∼Ub^A^ structure, both, WT UBR5 and UBR5^Dimer^ modify all possible di-UBs in vitro, with a preference for K11-linked chains.

## DISCUSSION

Our collection of cryo-EM reconstructions reveals the structural basis for linkage-specific Ub chain formation by a HECT E3, and together with published work defines a conserved step-by-step conformational trajectory for HECT E3-mediated Ub transfer cascades. To date, progression across an entire ubiquitylation cascade has not been published for any other HECT E3, however, a recent report of the yeast Ufd4 HECT E3 agrees with our proposed mechanism ^63^.Crystal structures of HECT domains from four different E3s – human NEDD4L, NEDD4, and HUWE1 and yeast Rsp5 – representing different states superimpose on our UBR5 structures and show HECT E3 ubiquitylation proceeds like a relay (Extended Data Figure 10a-d)^32–35^. These superimpose on the various UBR5 structures, and show HECT E3 ubiquitylation proceeds like a relay (Extended Data Figure 9a-d). First, as seen for UBR5 and at high-resolution for NEDD4L^32^, HECT domains receive Ub^D^ from E2 in the Inverted-T-configuration. This arrangement (1) co-positions the HECT domain N- and C-lobes and the E2∼Ub^D^ conjugate, (2) extends the E2∼Ub thioester bond, and (3) aligns the E2 and E3 active sites for Ub^D^ transfer between them^32^ (Extended Data Figure 9a). All the structures further suggest that HECT E3 C-lobes linked to Ub^D^ form a structural unit^32–35, 64^. Second, after formation of the E3∼Ub^D^ conjugate, this unit swivels around the N-lobe to achieve the L-configuration (Extended Data Figure 9b-c). Data for Rsp5^33^ and UBR5 indicate that the L-configuration transfers the Ub^D^ to a substrate, or to Ub^A^ during polyubiquitylation (Extended Data Figure 9d). We propose that L-shaped HECT∼Ub^D^ arrangements serve as platforms luring acceptors for further modification. Notably, the HUWE1 HECT domain-linked Ub^D^ complex^35^ not only overlays with UBR5^Dimer^∼Ub^D^, its C-terminal tail is configured as in the structure showing UBR5-mediated Ub chain formation. Our data suggest the HECT E3 C-tail arrangement allows penultimate hydrophobic side-chains to serve as molecular glues between the donor and acceptor Ubs, at least during K48-linked Ub chain formation.

Yet, UBR5’s catalytic HECT domain does not mediate polyubiquitylation on its own. This requires the acceptor Ub-binding UBA domain, and is substantially potentiated by full-length UBR5 (Figures 1i, 4c). UBR5 scaffold elements shape the Inverted-T TS1 and L-shaped TS2 catalytic configurations by unique positive and negative interactions (Figure 6). The L-arrangement is stabilized by UBR5’s HD and DSD domains binding on one side to the scaffold, and on the other to the HECT domain N- and C-lobes, respectively. Although E2 binding is incompatible with this arrangement between the scaffold and HECT domain, N-lobe binding to E2 in the alternative TS1 configuration is guided by simultaneous C-lobe binding to the E2’s linked Ub^D^. As such, severance of the E2∼Ub^D^ bond upon formation of the UBR5∼Ub^D^ intermediate re-enables the L-configuration, which is further stabilized by additional interactions from the HECT C-terminal tail and covalently-linked Ub^D^. Ub^A^ is recruited by the UBA domain in a manner compatible with a distal Ub in any chain type and more proximal Ub in some, while its acceptor residue 48 is guided into the active site by Ub^A^ interactions with the HECT domain N- and C-lobes including with the LOL, the C-terminal tail, and with Ub^D^. After Ub^D^ transfer to Ub^A^, affinity of the produced chain for UBR5 would be diminished by loss of interactions mediated directly by Ub^D^ linkage to UBR5, and loss of contacts upon dissolution of the E3 C-tail structure that also depends on UBR5 covalent linkage to Ub^D^. This would reset UBR5 for another round of Ub chain formation. Thus, the conformational trajectory forging linkage-specific Ub chains is achieved by the E2∼Ub^D^ and Ub^A^ substrates, and the reaction products, positively and negatively synergizing with numerous UBR5 structural features in a feed-forward manner.

**Figure 6:**
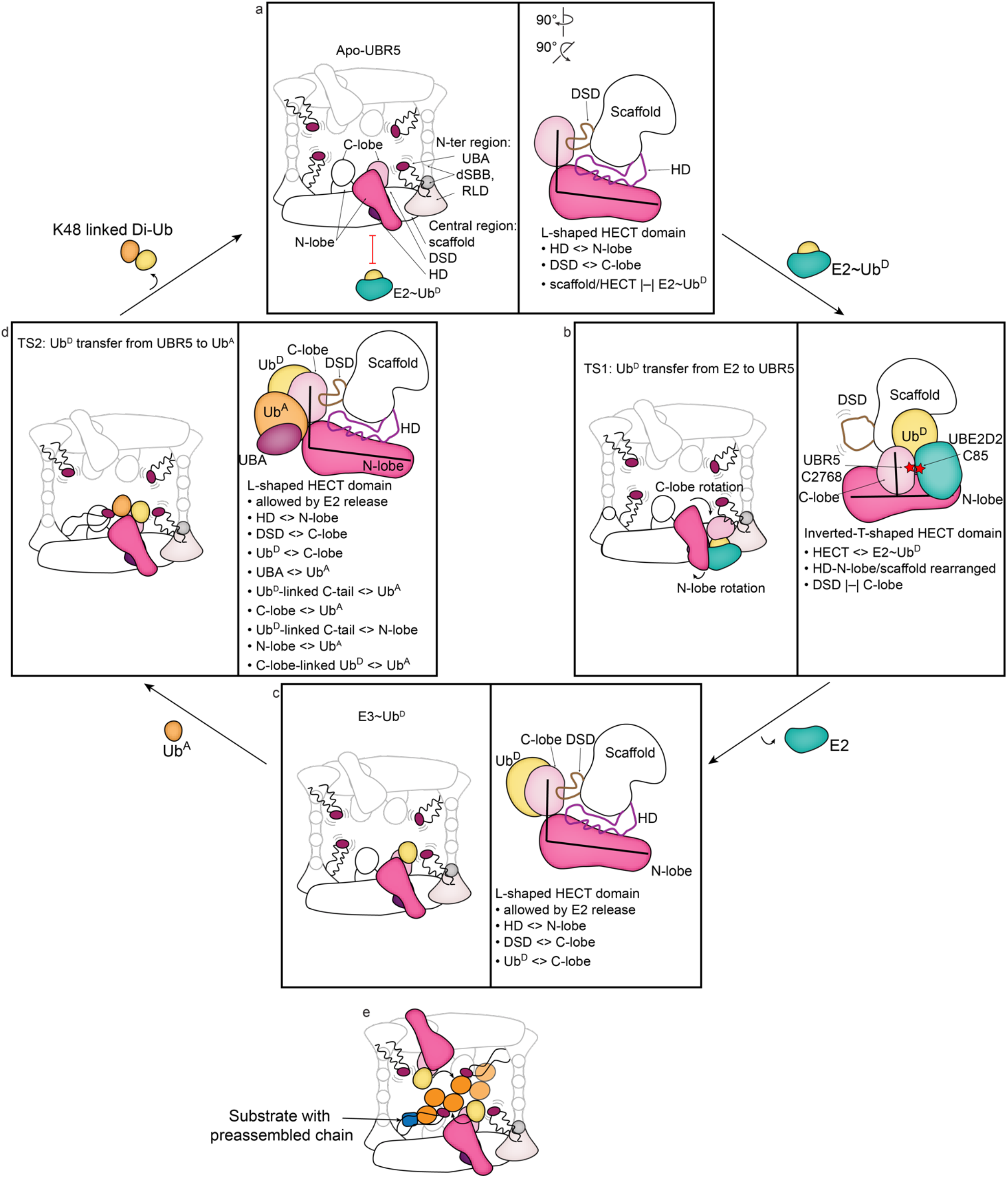
Feed-forward mechanism of K48-linked Ub chain formation by UBR5. **a,** The 2799-residue apo UBR5 forms a dimeric, U-shaped multidomain scaffold, which flexibly tethers Ub^A^-binding UBA domains, connects to the C-terminal HECT domains, and can further oligomerize into an oval-shaped tetramer. From the perspective of one HECT domain, apo UBR5 is poised in the L-configuration. Extensive interactions betwen UBR5 scaffold elements and HECT domain include “HECT Display” (HD) and Domain-Swapped-Dimerization (DSD) domains binding both the scaffold and HECT domain N- and C-lobes respectively. The scaffold/HECT domain arrangement, which is incompatible with binding E2∼Ub^D^, is relatively destablized compared to the TS2 assembly that is further supported by UBR5 covalent linkage to Ub^D^ and engagement of Ub^A^. **b,** Avid binding between E2 to HECT domain N-lobe and its covalently-linked Ub^D^ to the C-lobe drives scaffold/HECT reorientation, and supports the HECT domain Inverted-T-configuration juxtaposing E2 and E3 active sites for Ub^D^ transfer between them. **c,** After formation of UBR5∼Ub^D^ intermediate, release of free E2 would allow re-establishing scaffold connections with the HECT domain in the L-configuration. **d,** The acceptor Ub’s K48 and the UBR5-linked donor Ub’s C-terminus are juxtaposed through an extraordinary array of inter- and intra-protein interactions, depending on numerous UBR5 regions – the UBA domain, the scaffold, and the HECT domain – and between the two ubiquitins being adjoined. **e,** All pre-existing chain types – and especially K11-linked chains – could provide one or more ubiquitins structurally competent for further modification by UBR5. UBR5 oligomerization would allow avid binding and simultaneous modification of multiple ubiquitin moieties within a single chain or linked to various sites on a single substrate.

It seems likely that interactions between N-terminal elements and the C-terminal HECT domain establish functions across the E3 family. Interestingly, the only other HECT E3 for which full-length structures are available, HUWE1, also shows its many N-terminal substrate-binding domains arranged in a scaffold interacting with a peripherally-perched HECT domain^65, 66^. However, HUWE1’s HECT domain is maintained in the autoinhibited conformation. Moreover, structures have also shown N-terminal regions autoinhibiting NEDD4-family HECT E3s^67–69^. Thus, the overall UBR5 architecture stands out for conformationally priming the HECT domain for activity toward pre-ubiquitylated substrates. Although future studies will be required to visualize how post-translational modifications and substrates further modulate HECT E3 catalytic architectures^31, 36, 70, 71^, it seems that UBR5’s UBR and MLLE domains – facing and flexibly tethered to the HECT domain – are well situated to position recruited proteins adjacent to the L-shaped UBR5∼Ub^D^ active site for modification (Extended Data Figure 2b).

Finally, UBR5’s linkage-specific Ub chain-forming catalytic architecture is established in the context of an oligomer (Figure 1). Interestingly, a different E3 ligase, the RING-family GID complex, was also shown form an oligomer that determines E3 ligase function^72, 73^. GID E3 oligomerization allows: (1) multiple substrate binding domains to simultaneously engage degrons from multiple protomers of an oligomeric metabolic enzyme substrate; (2) multiple E3 ligase active sites to simultaneously ubiquitylate multiple lysines across the substrate complex; and (3) directing Ub to specific sites influencing regulation of the substrate^72^. For UBR5, oligomerization also provides multiple acceptor Ub-binding UBA domains, and multiple ubiquitylation active sites. The long flexible tethers between the RLD and UBA domains likely offer innumerable paths for avidly capturing multiple Ubs, linked to different sites on a substrate, and/or within pre-formed chains, for their K48 modification from the multiple HECT active sites (Figure 6e). Thus, taken together, the structural mechanisms explain why UBR5 is a highly efficient K48-linked Ub chain forming machine.

## ACKNOWLEDGEMENTS

We thank: D. Bollschweiler and T. Schäfer for assistance with cryo-EM; S. von Gronau for baculovirus and insect cell culture; S. Uebel for peptide-synthesis, the Schulman-department for discussions, especially J. Kellermann, J. Liwocha, D. Sherpa, L. Hopf, K. Baek, and J. Farnung. This study was supported by the Max Planck Gesellschaft, the ERC (H2020 789016-NEDD8Activate), and Leibniz Prize from the DFG (SCHU 3196/1-1).

## AUTHOR CONTRIBUTIONS

Biochemistry: L.A.H. CryoEM: L.A.H., D.H.-G., and J.R.P.. Structure building and refinement: J.R.P., L.A.H., and B.A.S.. Protein purification: L.A.H, D.H.-G., and R.V.. Reagent quality control design and contributed by D.T.V.. Preparation of chemically-modified ubiquitins: L.A.H., D.H.-G., D.A.P.B., M.P.C.M., and G.J.v.d.H.v.N.. Manuscript preparation: L.A.H. and B.A.S, with input from all authors.

## COMPETING INTEREST STATEMENT

BAS is a member of the scientific advisory boards of Interline Therapeutics and BioTheryX, and co-inventor of intellectual property licensed to Cinsano.

## DATA AVAILABILITY STATEMENT

The structural data will be available from EMDB and RCSB upon manuscript publication (UBR5^Dimer^ EMDB-16355, PDB 8C06; UBR5^Dimer^∼Ub^D^∼Ub^A^ EMDB-16356, PDB 8C07. Cryo-EM maps were deposited and are available via the following accession number: UBR5^C2768A^ EMDB-16865, UBR5^Dimer^∼Ub^D^∼UBE2D2 EMDB-16867, UBR5∼Ub^D^ EMDB-16866.). Raw gel images will be submitted upon manuscript revision, to be made available upon publication.

## Extended Data

**Extended Data Table 1:** Cryo-EM data collection, refinement, and validation statistics.

|  | Apo<br>Tetramer<br>(EMDB-16865) | Apo Dimer<br>(EMDB-16355)<br>(PDB<br>8C06) | E2-Ub-bound<br>Dimer<br>(EMDB-16867) | UbVME-bound<br>Dimer<br>(EMDB-16866) | K48-DiUb-<br>bound Dimer<br>(EMDB-16356)<br>(PDB 8C07) |
| --- | --- | --- | --- | --- | --- |
| **Data collection and processing** |  |  |  |  |  |
| Magnification | 64,000 | 105,000 | 73,000 | 22,000 | 105,000 |
| Voltage (kV) | 300 | 300 | 200 | 200 | 300 |
| Electron exposure<br>(e-/Å<sup>2</sup>) | 56.3 | 67.8 | 70 | 60 | 69 |
| Defocus range (μm) | -2.4 - -0.8 | -3.0- -0.5 | -3.5- -1.0 | -2.6- -<br>0.8 | -2.2 - -0.6 |
| Pixel size (Å) | 1.384 | 0.8512 | 1.997 | 1.885 | 0.8512 |
| Symmetry imposed | C2 | C2 | C1 | C1 | C1 |
| Initial particle<br>images (no.) | 1,122,292 | 762,722 | 1,381,245 | 834,722 | 1,708,682 |
| Final particle<br>images (no.) | 148,825 | 226,919 | 46,615 | 197,281 | 141,034 |
| Map resolution (Å) | 3.7 | 2.7 | 7.3 | 5.3 | 3.3 |
| FSC threshold | (0.143) | (0.143) | (0.143) | (0.143) | (0.143) |
| Map resolution<br>range (Å) | 3.3-9.8 | 2.6-5.1 | 5.6-25.1 | 4.7-18.2 | 3.2-10 |
| **Refinement** |  |  |  |  |  |
| Initial model used<br>(PDB code) |  | AlphaFold2 |  |  | 8C06 |
| Model resolution<br>(Å) |  | 2.7 |  |  | 3.3 |
| Model composition |  |  |  |  |  |
| Non-hydrogen<br>atoms |  | 25332<br>3298 |  |  | 4974<br>644 |
| Protein residues |  | 6 |  |  | 1 |
| Ligands |  |  |  |  |  |
| *B* factors (Å<sup>2</sup>) |  |  |  |  |  |
| Protein |  | 62 |  |  | 81.2 |
| Ligand |  | 52 |  |  | 84 |
| R.m.s. deviations |  |  |  |  |  |
| Bond lengths (Å) |  | 0.003 |  |  | 0.004 |
| Bond angles (°) |  | 0.765 |  |  | 0.944 |
| Validation |  |  |  |  |  |
| MolProbity score |  | 1.4 |  |  | 1.7 |
| Clashscore |  | 2.3 |  |  | 4.1 |
| Poor rotamers<br>(%) |  | 0 |  |  | 0 |
| Ramachandran<br>plot |  | 95 |  |  | 92 |
| Favored (%) |  | 5 |  |  | 8 |
| Allowed (%) |  | 0 |  |  | 0 |
| Disallowed (%) |  |  |  |  |  |

**Extended Data Figure 1:**
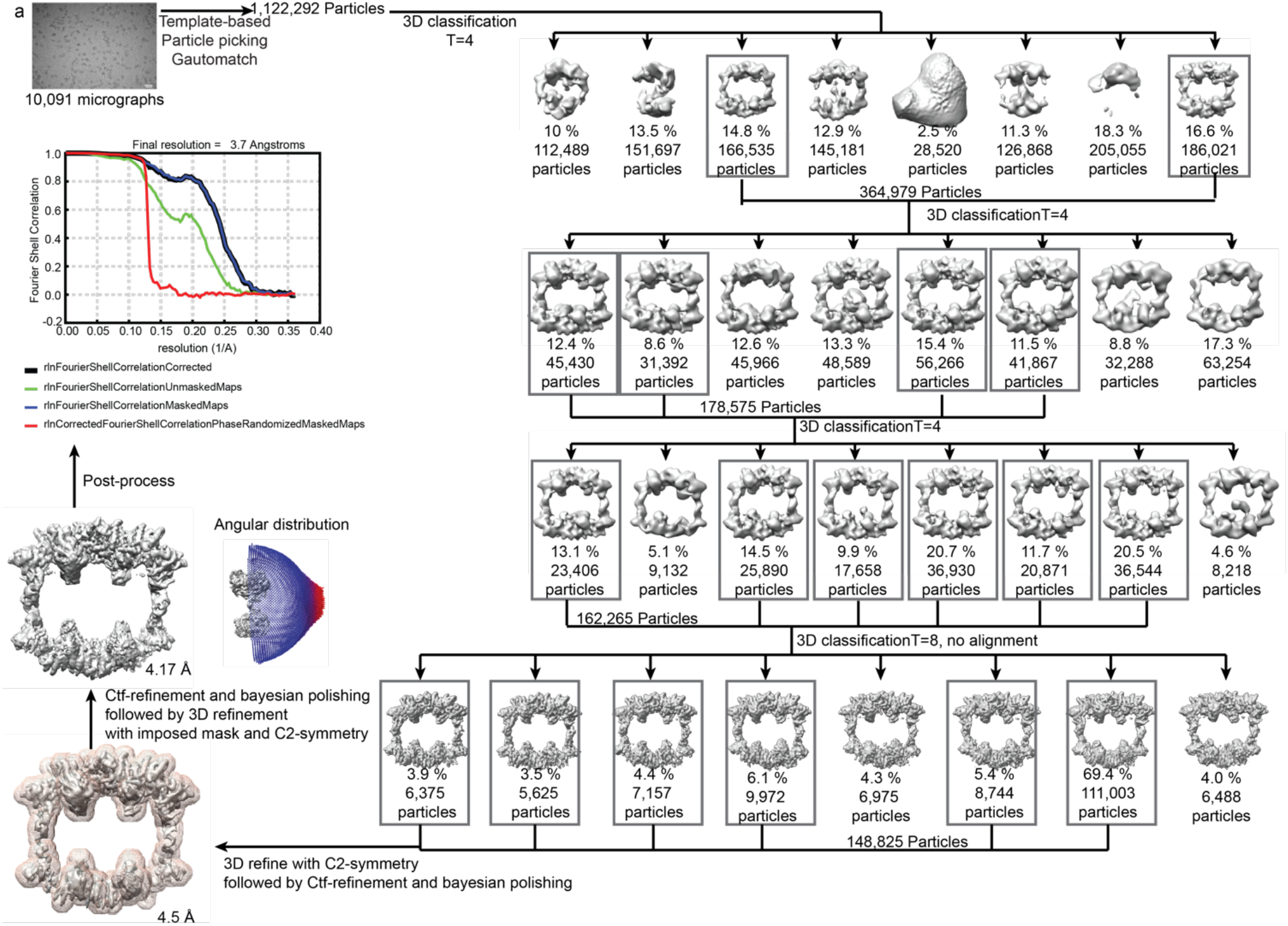
Cryo-EM processing scheme of UBR5^C2768A^. Cryo-EM processing schematic of UBR5^C2768A^. Scalebar on micrograph corresponds to 500 Å. Data processed in RELION 3.1.1 yielded a 3D reconstruction with a resolution of 3.7 Å by the gold-standard Fourier shell correlation of 0.143.

**Extended Data Figure 2:**
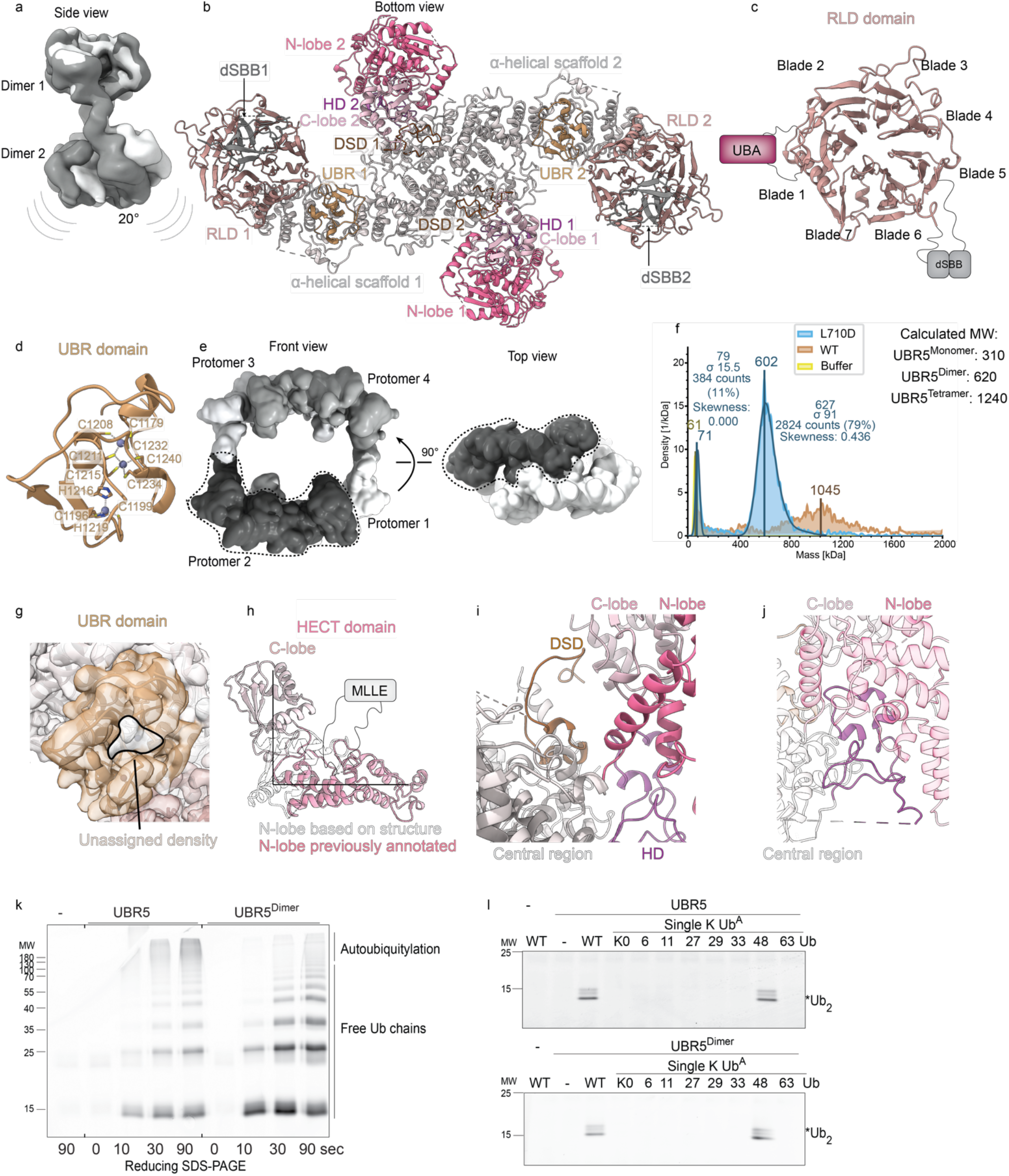
Structural and functional characterization of UBR5 and UBR5^Dimer^. **a,** Superposition of one U-shaped dimer within different 3D classes of UBR5^C2768A^ processing shows differences in orientations of U-shaped dimers within tetrameric WT UBR5 assembly. The two maps are shown in the side view, one in grey and the other white. **b**, Bottom-view of UBR5^Dimer^, annotating the RLD, UBR, ⍺-helical scaffold, DSD, HD and HECT domain N- and C-lobes, indicating protomers 1 or 2 of the dimer. The dSBB domains are only poorly visible in the map, but one of the two barrels could be placed in the density and is indicated. **c,** Structure of the RLD. Each UBR5 protomer contains a previously un-annotated discontinuous 7-bladed beta-propeller resembling the RCC1-like domain (RLD) found in HERC-family HECT E3 ligases. The first blade initiates at UBR5’s N-terminus, and is completed by two beta-strands from the C-terminal portion of the beta-propeller. The second blade harbors three antiparallel strands followed by a ∼270 residue insertion that includes the UBA domain (purple bar). The insertion is not visible in the cryo-EM density for UBR5^Dimer^, and is followed by the fourth strand of blade-2. The third to fifth blades are conventional, but the loop connecting the fifth and the sixth blades is interrupted by a >220-residue long insertion that contains the dSBB domain mediating interactions between the two U-shaped dimers in the tetramer. **d,** The zinc-binding UBR domain is positioned within the UBR5 scaffold, facing the opposite HECT domain in trans. Cys1196, Cys1199, His1216, and His1219 coordinate one zinc ion. Another zinc is coordinated by Cys1215, Cys1234, Cys1240, and Cys1211, which is shared with the third zinc ion, coordinated additionally by Cys1179, Cys1208, and Cys1232. **e,** Protomers assigned based on experimentally-derived UBR5^Dimer^ structure and AlphaFold2 predictions in tetramer are shown the front and top views. The four monomers within a tetramer are shaded individually and one monomer is further highlighted using a dotted line. **f,** Molecular weights of UBR5^WT^ and UBR5^L710D^ assemblies as determined by mass photometry. The main peaks for the respective samples are highlighted, along with measured and calculated MWs for a dimer and a tetramer. **g,** Cryo-EM map over the UBR domain shows unassigned density in the canonical peptide– binding cleft. **h,** L-shaped HECT domain from structure of UBR5^Dimer^, in white. The previously annotated N-lobe is shown in dark pink and C-lobe in light pink. **i,** Close-up of the domain-swapped-dimerization domain (DSD, residues 1691-1720, brown). The DSD meanders between the scaffold and HECT domain C-lobe. **J,** Zoom-in of HECT Display domain (HD, residues 2016-2076), which both interacts with the central region and nestles in the HECT domain N-lobe. The HD domain binds the opposite side of the N-lobe from the canonical E2-binding site. It also contacts the C-lobe and the interlobe linker connecting the N- and C-lobes. **k,** Autoubiquitylation/free Ub chain formation in pulse-chase format for UBR5 and UBR5^Dimer^. Fluorescent (*) Ub was used and detected in reducing SDS-PAGE gels. **l,** Di-Ub synthesis assay testing linkage-specificity of WT UBR5 and UBR5^Dimer^. UBE2D2∼*Ub was added to the indicated versions of Ub^A^ and either WT UBR5 or UBR5^Dimer^. Generation of the di-Ub product was analyzed by fluorescent scanning after reducing SDS-PAGE. K0=all lysines mutated to arginine. The other Ubs are indicated by their sole lysine, with all other lysines mutated to arginines.

**Extended Data Figure 3:**
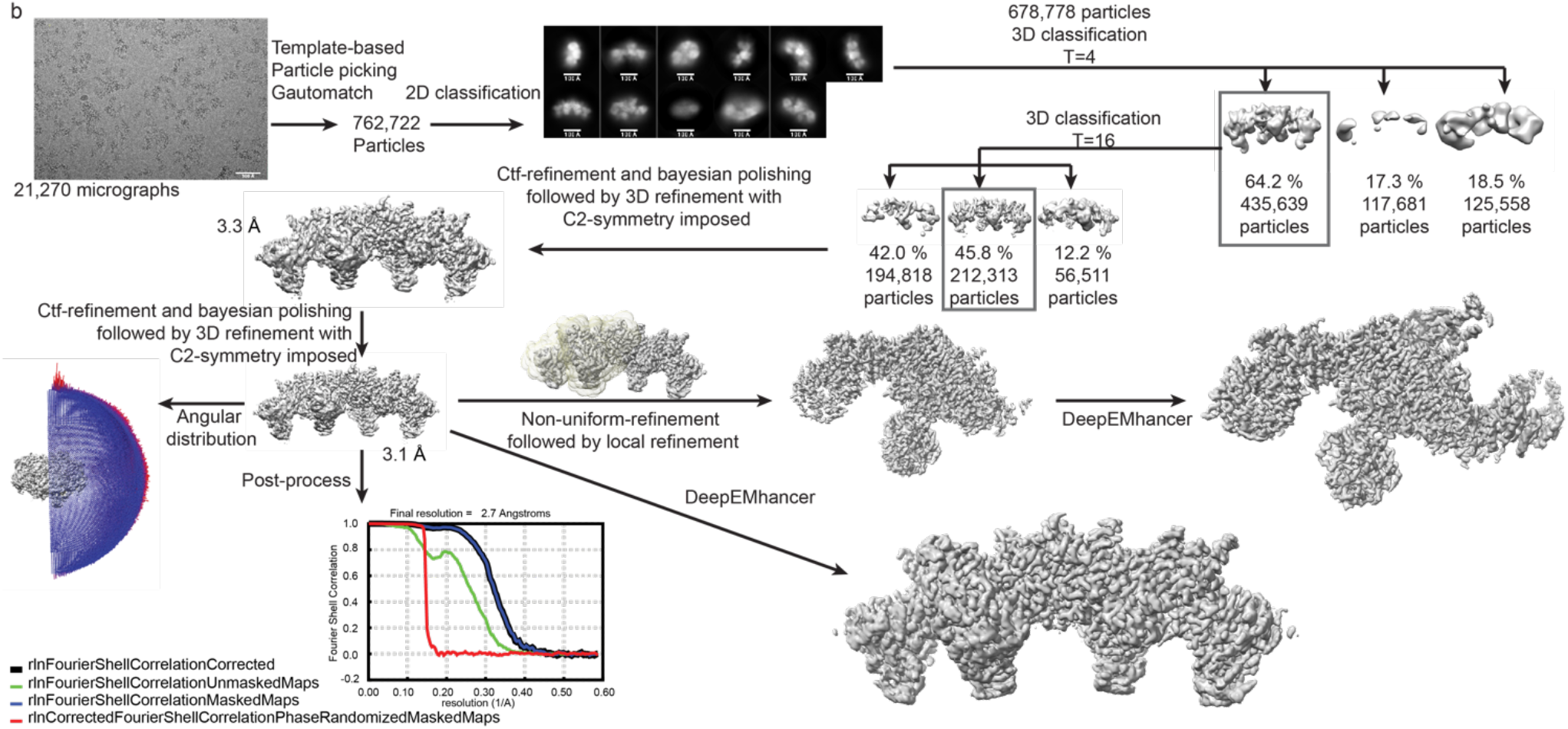
Cryo-EM processing scheme of UBR5^Dimer^. Cryo-EM processing scheme of UBR5^Dimer^, with the L710D mutation. Scalebar on micrograph corresponds to 500 Å. Scalebars on 2D classes correspond to 100Å. Mask imposed for local refinement subsequently to unmasked non-uniform refinement is shown in yellow transparent surface. Data processing was performed in RELION 4.0 followed by processing in CryoSparc4.2.0 for non-uniform refinement and local refinement. A final 3D reconstruction with a resolution of 2.7 Å by the gold-standard Fourier shell correlation of 0.143 and a local refined reconstruction of 3 Å was achieved.

**Extended Data Figure 4:**
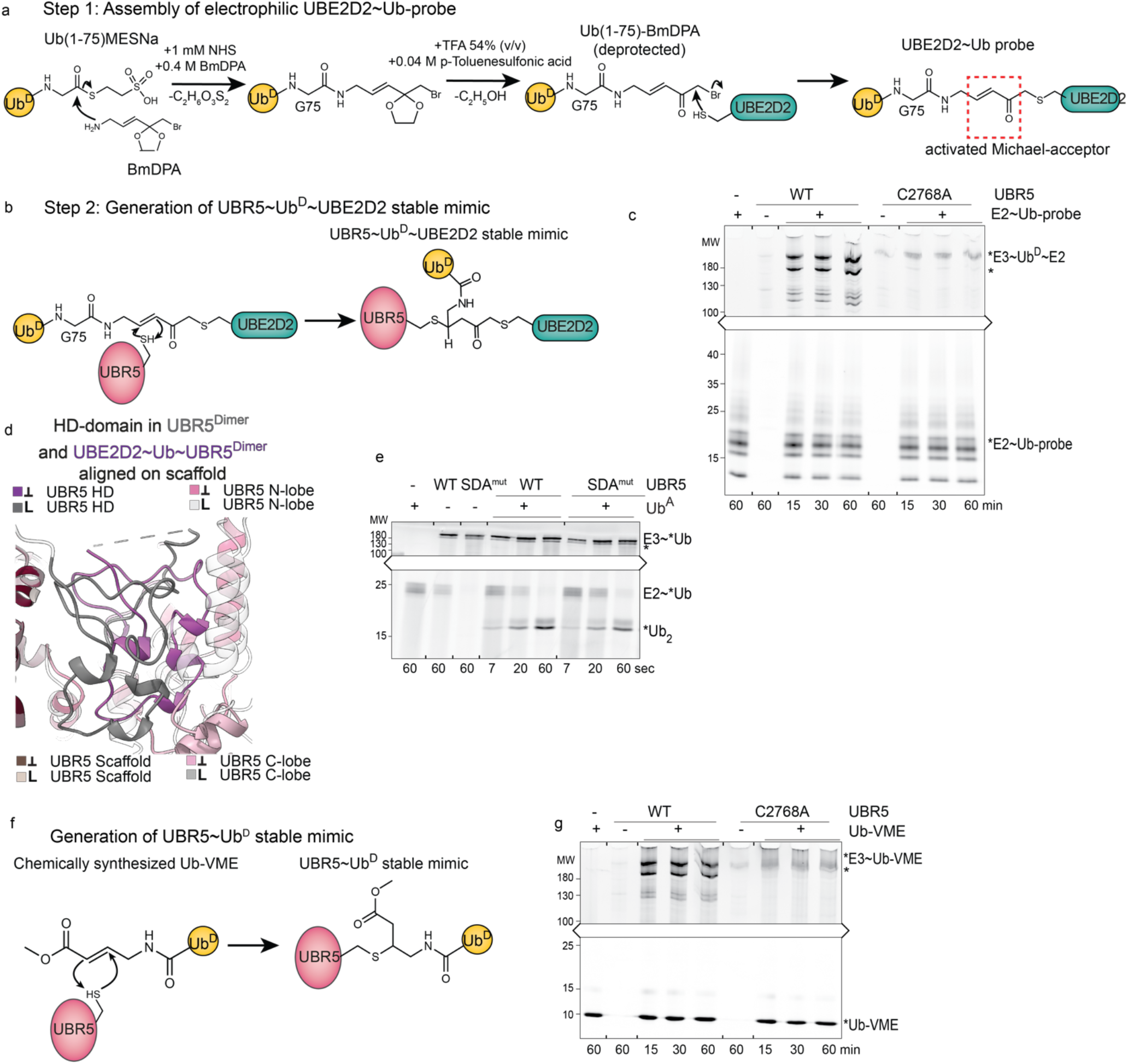
Chemical biology tools to visualize Ub^D^ transfer from E2 to UBR5 and subsequent UBR5∼Ub^D^ intermediate. **a,** Stable mimic representing TS1, Ub^D^ transfer from an E2 to UBR5 was generated in two steps. Reaction scheme for the first step, whereby an electrophile was installed between Ub^D^ C-terminus (here, Ub^D^’s C-terminal G76 is deleted to approach near-native geometry) and active site Cys of E2 UBE2D2 (with other cysteines mutated as described in Methods). **b,** Reaction scheme for second step in generation of TS1 mimic. The UBE2D2∼Ub^D^ probe was reacted with UBR5^Dimer^ to generate a UBR5^Dimer^∼Ub^D^∼UBE2D2 complex wherein the E2 UBE2D2, Ub’s C-terminus, and UBR5^Dimer^ are linked at a single atom. **c,** UBR5 catalytic Cys dependence for forming the UBR5^Dimer^∼Ub^D^∼UBE2D2 complex was tested using a version of the E2∼Ub^D^ probe with fluorescent UBE2D2. Reactivity with UBR5 was then analyzed by fluorescent scanning after SDS-PAGE for reaction products. Again, the faster-migrating band, marked *, is a degradation-product of UBR5 that could not be separated from full-length protein during purification. **d,** Close-up comparing HD domain orientations in apo UBR5^Dimer^ and UBR5^Dimer^∼Ub^D^∼UBE2D2, in grey and purple, respectively. Structure of apo UBR5^Dimer^ and model of UBR5^Dimer^∼Ub^D^∼UBE2D2 were overlaid over the scaffold, and domains contacting the HD domain are also shown. The colored squares indicate the L- and inverted-T-conformations of the HECT domain in the two complexes, and the shading of the scaffold, HD domain, and HECT domain N- and C-lobes in apo UBR5^Dimer^ and UBR5^Dimer^∼Ub^D^∼UBE2D2, respectively. **e,** Di-Ub-synthesis assay for the SDA mutant. **f,** Reaction scheme to generate stable mimic of UBR5∼Ub^D^. Fully synthetic Ub-VME was reacted with UBR5^Dimer^. Trap specificity of Ub-VME towards catalytic cysteine of UBR5. UBR5 catalytic Cys dependence for forming the UBR5∼Ub^D^ complex was tested using a fluorescent version of Ub-VME. Reactivity with UBR5 was then analyzed by fluorescent scanning after SDS-PAGE for reaction products.

**Extended Data Figure 5:**
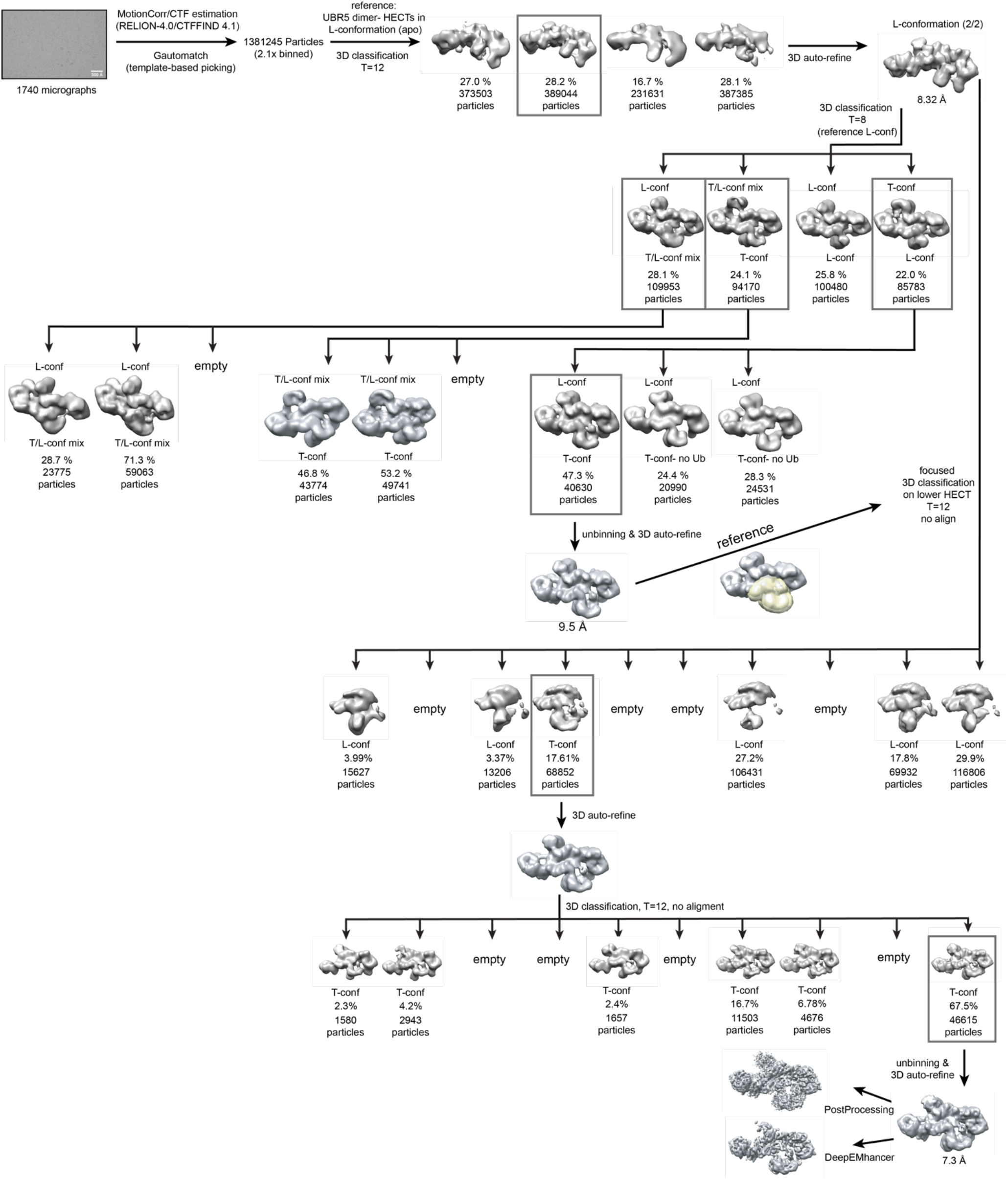
Cryo-EM processing scheme of stable mimic representing UBR5∼Ub^D^∼E2. Cryo-EM processing schematic of UBR5^Dimer^∼Ub^D^∼UBE2D2. Scalebar on micrograph corresponds to 500 Å. Mask used for focused 3D-classification is shown in yellow transparent surface. Data processed in RELION 4.0 yielded a final 3D reconstruction with a resolution of 7.3 Å by the gold-standard Fourier shell correlation of 0.143.

**Extended Data Figure 6:**
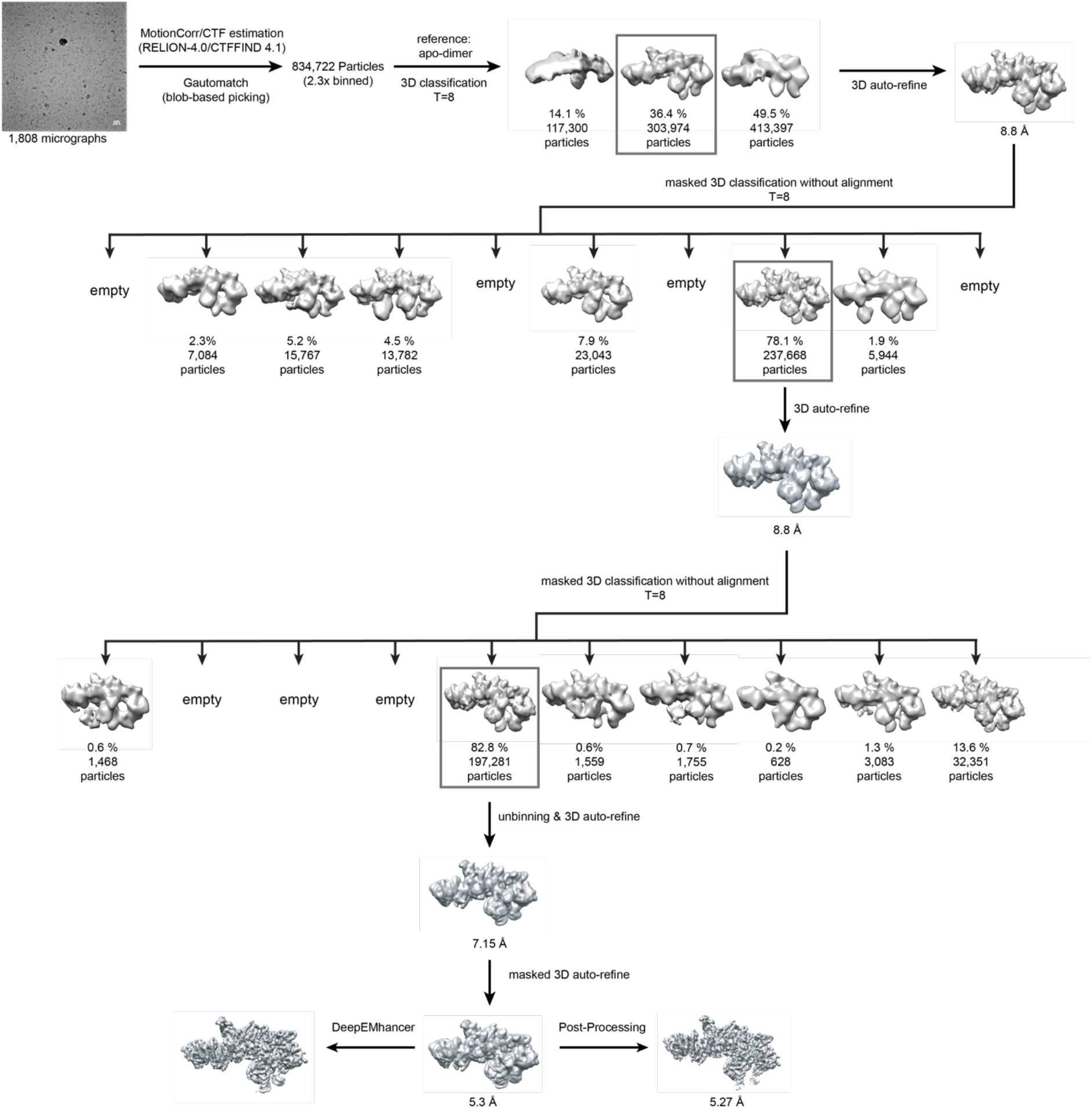
Cryo-EM processing scheme of stable mimic representing UBR5∼Ub^D^. Cryo-EM processing schematic of UBR5^Dimer^∼Ub^D^. A scalebar on the micrograph represents 300 Å. Data processing was performed using RELION 4.0 and yielded a final 3D reconstruction with a resolution of 5.3 Å by the gold-standard Fourier shell correlation of 0.143.

**Extended Data Figure 7:**
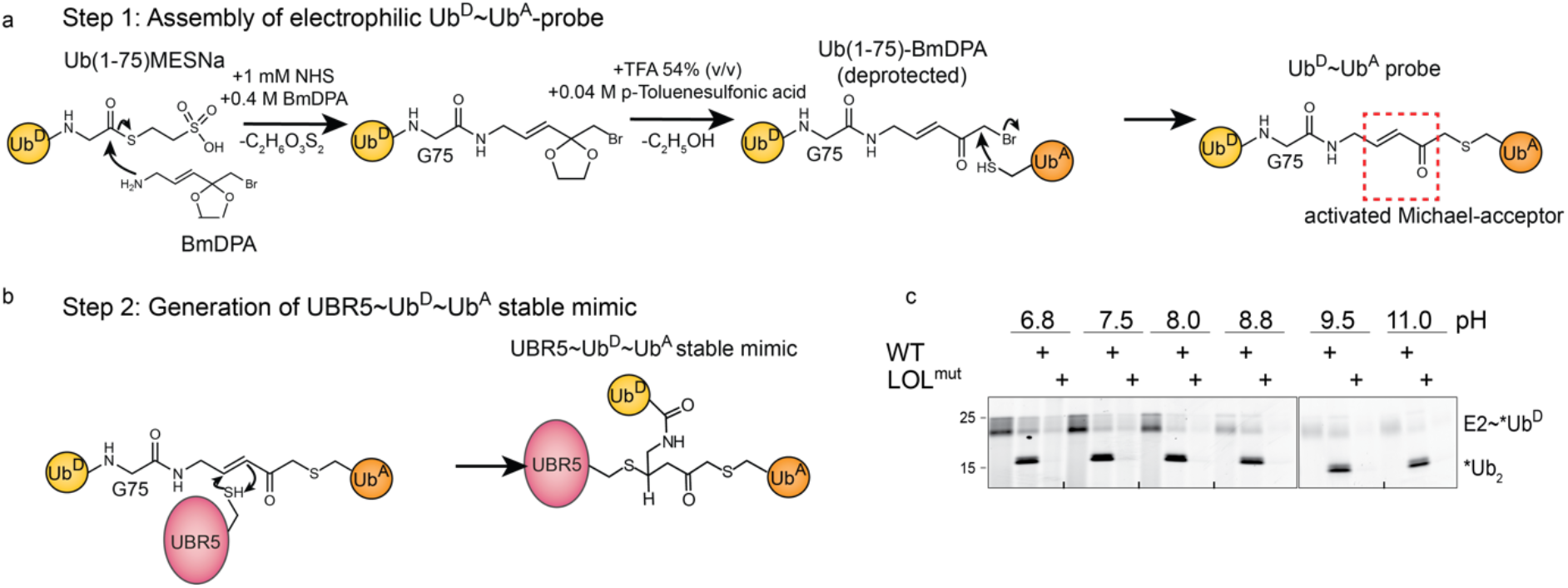
Chemical biology tools to visualize K48-linked Ub chain formation by UBR5. **a,** Stable mimic representing TS2, Ub^D^ transfer from UBR5 to an Ub^A^, was generated in two steps. Reaction scheme for the first step, whereby an electrophile was installed between Ub^D^ C-terminus (here, Ub^D^’s C-terminal G76 is deleted to approach near-native geometry) and a Cys replacement for K48 in Ub^A^. **b,** Reaction scheme for second step in generation of TS2 mimic. The Ub^D^∼Ub^A^ probe was reacted with UBR5^Dimer^ to generate a UBR5^Dimer^∼Ub^D^∼Ub^A^ complex wherein the C-terminus of the Ub representing the donor, residue 48 of the Ub representing the acceptor, and a HECT domain catalytic Cys in UBR5^Dimer^ are linked at a single atom. **c,** Di-Ub synthesis assay testing if the deficiency in di-Ub synthesis caused by LOL mutations is pH dependent. UBE2D2∼*Ub^D^ was added to WT or LOL mutant UBR5 at the indicated pH. Reaction products were separated by non-reducing SDS-PAGE and imaged by fluorescent scanning.

**Extended Data Figure 8:**
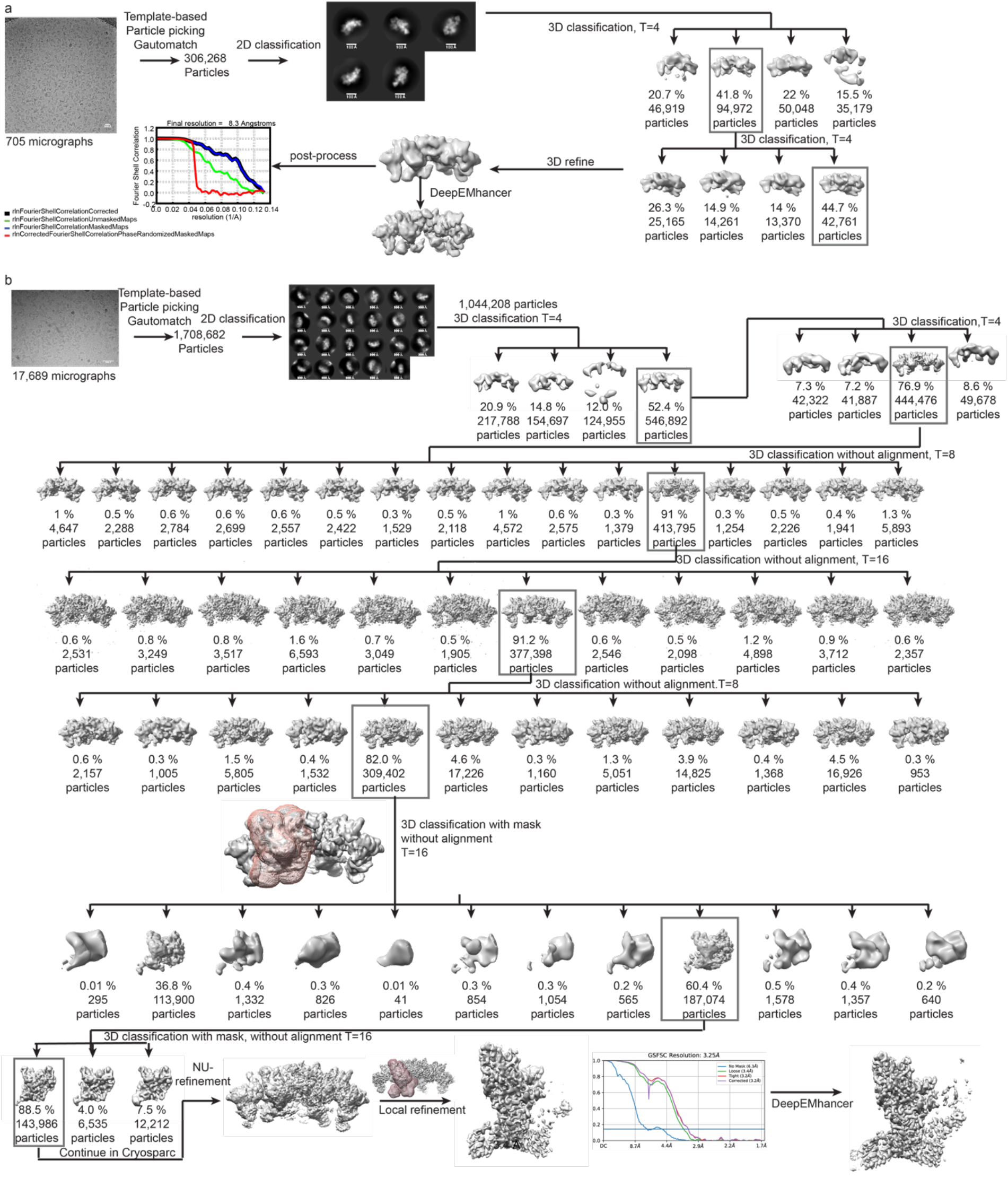
Cryo-EM processing scheme of stable mimic representing K48-linked Ub chain formation by UBR5. **a,** Cryo-EM processing schematic of UBR5^Dimer^∼Ub^D^∼Ub^A^ complex for overall map. Scalebar on micrograph corresponds to 300 Å. Scalebars on 2D classes correspond to 100Å. Data processing was performed using RELION 3.1.1 and yielded a final 3D reconstruction with a resolution of 8.3 Å by the gold-standard Fourier shell correlation of 0.143 **b,** Cryo-EM processing schematic of UBR5^Dimer^∼Ub^D^∼Ub^A^ complex. Scalebar on micrograph corresponds to 500 Å. Scalebars on 2D classes correspond to 100Å. Masks used at various steps of focused 3D classification and 3D refinement are shown in pink transparent surfaces. Data processing was performed using RELION 3.1.1 until focused 3D classification. Subsequent non-uniform refinement and focused 3D refinement were performed using Cryosparc yielded a final 3D reconstruction with a resolution of 3.4 Å by the gold-standard Fourier shell correlation of 0.143.

**Extended Data Figure 9:**
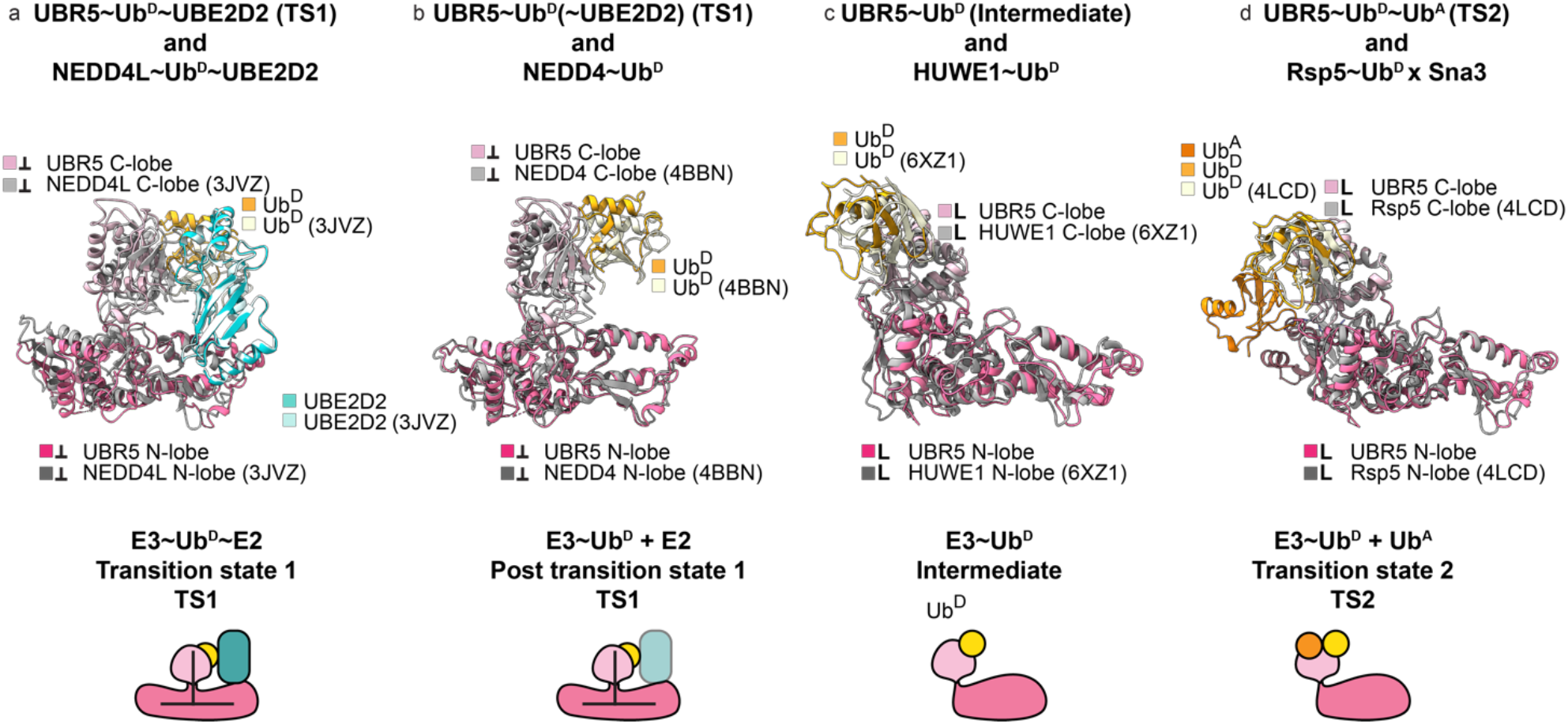
Conserved step-by-step conformational trajectory for HECT E3-catalyzed Ub transfer cascades. **a,** Structural superposition of UBE2D2∼UbD-bound HECT domains of UBR5 and NEDD4L (PDB: 3JVZ). The structures suggest E2 transfers Ub^D^ to HECT domains in the Inverted-T-configuration. Colored boxes indicate the respective complex as well as the conformation of the HECT domain in this complex. **b,** Structural superposition of Ub^D^ linked to HECT domain of UBR5 (TS1 model) and NEDD4 (PDB: 4BBN). This represents the Ub^D^-linked HECT domain in the Inverted-T-configuration, immediately after Ub^D^ transfer from E2. **c,** Structural superposition of Ub^D^ linked to HECT domains of UBR5 (from UBR5^Dimer^∼Ub^D^ complex) and HUWE1∼Ub^D^ (PDB: 6XZ1). Both show the HECT domain in the L-configuration. **d,** Structural superposition of HECT domains representing ubiquitylation complexes, K48-linked Ub chain formation by UBR5, and Ub^D^ transfer to a peptide substrate for Rsp5 (PDB: 4LCD). The structures suggest HECT domains transfer Ub^D^ to downstream targets – either substrates or acceptor Ub during polyubiquitylation – in the L-configuration.

## METHODS

### Construct design

All constructs in this study were generated using standard molecular biology techniques and verified using Sanger sequencing. The following constructs were used for protein expression:

| Protein | Construct | Vector | Expression system |
| --- | --- | --- | --- |
| E1 | GST-TEV-UBA1 | pLIB | High-Five insect cells |
| E2 | GST-TEV-UBE2D2 | pGEX | BL21(DE3) RIL cells |
|  | GST-TEV-UBE2D3 | pGEX | BL21(DE3) RIL cells |
|  | GST-TEV-UBE2D3<sup>F62A</sup> | pGEX | BL21(DE3) RIL cells |
| Donor ubiquitin | GST-3C-Cys-Ub | pGEX | BL21(DE3) RIL cells |
|  | GST-3C-Cys-Ub<sup>K0</sup> | pGEX | BL21(DE3) RIL cells |
|  | GST-3C-Cys-Ub<sup>K48R</sup> | pGEX | BL21(DE3) RIL cells |
| Single K ubiquitin | GST-3C-Ub | pGEX | BL21(DE3) RIL cells |
|  | GST-3C-Ub<sup>K0</sup> | pGEX | BL21(DE3) RIL cells |
|  | GST-3C-Ub<sup>K6only</sup> | pGEX | BL21(DE3) RIL cells |
|  | GST-3C-Ub<sup>K11only</sup> | pGEX | BL21(DE3) RIL cells |
|  | GST-3C-Ub<sup>K27only</sup> | pGEX | BL21(DE3) RIL cells |
|  | GST-3C-Ub<sup>K29only</sup> | pGEX | BL21(DE3) RIL cells |
|  | GST-3C-Ub<sup>K33only</sup> | pGEX | BL21(DE3) RIL cells |
|  | GST-3C-Ub<sup>K48only</sup> | pGEX | BL21(DE3) RIL cells |
|  | GST-3C-Ub<sup>K63only</sup> | pGEX | BL21(DE3) RIL cells |
| M1 linked Di-Ub | GST-TEV-Ub<sub>2</sub> | pGEX | BL21(DE3) RIL cells |
| Acceptor ubiquitin | Ub(1-76)-6xHis | pRSF | BL21(DE3) RIL cells |
|  | Ub(1-76)<sup>L8A,T9A</sup>-6xHis | pRSF | BL21(DE3) RIL cells |
|  | Ub(1-76)<sup>I44A</sup>-6xHis | pRSF | BL21(DE3) RIL cells |
|  | Ub(1-76)<sup>V70A L71A</sup>-6xHis | pRSF | BL21(DE3) RIL cells |
|  | Ub(1-76)<sup>H68A</sup>-6xHis | pRSF | BL21(DE3) RIL cells |
|  | Ub(1-76)<sup>R54E</sup>-6xHis | pRSF | BL21(DE3) RIL cells |
|  | Ub(1-76)<sup>A46F</sup>-6xHis | pRSF | BL21(DE3) RIL cells |
|  | Ub(1-76)<sup>A46D</sup>-6xHis | pRSF | BL21(DE3) RIL cells |
|  | Ub(1-76)<sup>D58A</sup>-6xHis | pRSF | BL21(DE3) RIL cells |
|  | Ub(1-76)<sup>D58R</sup>-6xHis | pRSF | BL21(DE3) RIL cells |
| HECT domain | GST-TEV-HECT<sup>withMLLE</sup> | pFLN | High-Five insect cells |
|  | GST-TEV-HECT<sup>ΔMLLE</sup> | pFLN | High-Five insect cells |
|  | GST-TEV-UBA-(GSG)<sub>5</sub>-HECT | pFLN | High-Five insect cells |
| UBR5-mutants | TwinStrep-GFP-3C-UBR5 | pEG | HEK293S, BacMam-system |
|  | TwinStrep-GFP-3C-UBR5<sup>C2768A</sup> | pEG | HEK293S, BacMam-system |
|  | TwinStrep-GFP-3C-UBR5<sup>L710D</sup><br>(UBR5<sup>Dimer</sup>) | pEG | HEK293S, BacMam-system |
|  | TwinStrep-GFP-3C-UBR5<sup>HLL1362-1364DDD</sup><br>(SDA<sup>mut</sup>) | pEG | HEK293S, BacMam-system |
|  | TwinStrep-GFP-3C-UBR5<sup>A2790W</sup> | pEG | HEK293S, BacMam-system |
|  | TwinStrep-GFP-3C-UBR5<sup>L224D</sup> | pEG | HEK293S, BacMam-system |
|  | TwinStrep-GFP-3C-UBR5<sup>D2283-2287A</sup><br>(LOL<sup>mut</sup>) | pEG | HEK293S, BacMam-system |
|  | TwinStrep-GFP-3C-UBR5<sup>L710D,E2287A</sup> | pEG | HEK293S, BacMam-system |
|  | TwinStrep-GFP-3C-UBR5<sup>F2796A</sup> | pEG | HEK293S, BacMam-system |
|  | TwinStrep-GFP-3C-UBR5<sup>F2798A</sup> | pEG | HEK293S, BacMam-system |
|  | TwinStrep-GFP-3C-UBR5<sup>Δ2799</sup> | pEG | HEK293S, BacMam-system |
|  | TwinStrep-GFP-3C-UBR5<sup>L710D,Y2773F</sup> | pEG | HEK293S, BacMam-system |
| Probe-constructs | His-Ub(1-75)-Intein | pET | BL21(DE3) RIL cells |
|  | Ub<sup>K48C</sup> | pET | BL21(DE3) RIL cells |
|  | GST-TEV-UBE2D2<sup>C21A,C107A,C111S</sup> | pGEX | BL21(DE3) RIL cells |

### Protein expression and purification preparation

Expression of ubiquitin, ubiquitin mutants and UBE2D2 was performed in *E. coli* BL21(DE3) RIL cells. BL21(DE) RIL cells containing the respective plasmid, were grown at 37°C in TB medium, supplemented with appropriate antibiotics. Upon reaching an optical density of 0.8, temperature was lowered to 18°C and induction was induced by adding IPTG to a final concentration of 0.6 mM. Consequently, expression was conducted for 18 h, before harvesting cells by centrifugation at 4°C for 15 min, 4.500 x g. The pellet was resuspended in lysis buffer containing 50 mM Tris-HCl pH 8.0, 200 mM NaCl, 5 mM DTT or 5 mM β-mercaptoethanol for His-tagged constructs, 2.5 mM PMSF, and additionally 10 μg/mL leupeptin, 20 μg/mL aprotinin, and 10 μg/mL DNase for constructs expressed in insect cells of HEK293S-cells. Cells were lysed by sonication and lysate was pre-cleared by centrifugation for 40 min at 4°C at 20.000 x g.

#### His-tagged proteins

His-tagged acceptor ubiquitins were purified via Ni-NTA-affinity chromatography in a gravity flow column setup. The resin was washed with 20 mM imidazole, 50 mM Tris-HCl pH 8.0, 200 mM NaCl, and 5 mM β-mercaptoethanol. Elution of specifically bound proteins was achieved with 300 mM imidazole. Subsequently, size exclusion chromatography was performed to rebuffer into 25 mM HEPES pH 7.5, 150 mM NaCl and yielded pure His-tagged protein.

#### GST-tagged proteins

E2 enzymes, single lysine acceptor ubiquitins, and M1-linked Di-Ub were purified by incubating pre-cleared lysate with GST-sepharose resin for 1 h at 4°C. After extensively washing the resin with 10 column volumes of wash buffer (50 mM Tris-HCl pH 8.0, 200 mM NaCl, 5 mM DTT), GST-fusion protein was eluted using 10 mM reduced glutathione. Cleavage of the GST-tag was achieved via proteolytic digest overnight at 4°C by addition of either His-tagged PreScission protease (in-house) for single lysine ubiquitin-constructs or His-tagged TEV protease for E2 enzymes and M1-linked Di-Ub. E2 and Di-Ub protein was subsequently purified using ion-exchange (IEX) chromatography and size exclusion chromatography with 25 mM HEPES pH 7.5, 150 mM NaCl (+1 mM DTT for E2). The single lysine ubiquitins were further purified using size exclusion chromatography in a buffer containing 25 mM HEPES pH 7.5, 150 mM NaCl subsequent to affinity purification.

#### Fluorescently-labeled ubiquitin

Donor ubiquitins, which were ultimately fluorescently labeled, were expressed as GST-3C-fusions in a pGEX-vector with an additional N-terminal cysteine. The purification and labelling schemes were adapted^74^. In brief, GST-affinity chromatography followed by PreScission cleavage was performed. On the next day, size exclusion chromatography into 50 mM HEPES pH 7.5, 200 mM NaCl, 5 mM DTT was carried out. The high concentration of reducing reagent is crucial to ensure complete reduction of the N-terminal cysteine, which is modified later. Before coupling to maleimide, DTT was removed by desalting twice with Zeba^TM^ Spin desalting columns into reaction buffer (50 mM HEPES pH 7.5, 150 mM NaCl). Fluoresceine-5-maleimide or tetramethylrhodamine-5-Maleimide (TAMRA) were resuspended in anhydrous DMSO and added to desalted ubiquitin in a 10-fold molar excess while keeping the final DMSO concentration below 5 %. The reaction was incubated for 2 h at room temperature before being quenched by the addition of 10 mM DTT. Consequently, desalting of samples was repeated to remove unreacted maleimide, followed by two size exclusion chromatography runs to yield highly pure labeled ubiquitin conjugates. Di-Ub synthesis assays were performed with TAMRA-labeled Ub with Lys 48 mutated to Arg to prevent its use as acceptor. The autoubiquitylation assay to compare UBR5 and UBR5^Dimer^, as well as the polyubiquitylation assay to compare UBR5 with the SDA-mutant were performed using fluoresceine-labeled WT Ub.

#### Tagless ubiquitin via acidic precipitation

Tagless ubiquitins were used as basis for the generation of various ubiquitin chains, which were employed as acceptor chain in the Tri-Ub synthesis assay in Figure 6e, and added to improve cryo-EM samples. Additionally, Ub^K48C^ was one of the building blocks for formation of the Di-Ub-probe. To obtain tagless ubiquitin, it was expressed using a pET22b-vector. Expression as well as cell lysis was conducted as described earlier. Next, acetic acid was slowly added to the lysate until a pH of ∼4.5 was reached to precipitate out most other proteins except for ubiquitin. Subsequent ion exchange chromatography of the cleared supernatant, was followed by size exclusion chromatography into 25 mM HEPES pH 7.5, 150 mM NaCl (+ 1 mM DTT in case of Ub^K48C^) to yield the desired tagless ubiquitin and ubiquitin mutants.

#### Insect cell-derived proteins

With the correct sequence in hand, we pioneered the expression of the isolated UBR5 HECT domain, with or without interrupting MLLE domain, as well as a UBA-HECT-fusion, in Hi5 insect cells. These constructs contained an N-terminal GST-tag followed by a TEV-cleavage site. Additionally, human UBA1 was also expressed as GST-TEV-fusion in the same cell system. Cell lysis and pre-clearance of the lysate were performed as mentioned previously for bacterial expressions. Protein purification was performed using gravity flow affinity purification with GST-resin, followed by proteolytic cleavage of GST-fusion protein with His-tagged TEV-protease. Finally, ion exchange chromatography and size exclusion chromatography were carried out with the final buffer consisting of 25 mM HEPES pH 7.5, 150 mM NaCl, 1 mM DTT.

#### Human cell-derived UBR5

The GFP-UBR5 plasmid was a gift from Darren Saunders (Addgene plasmid #52050 ;http://n2t.net/addgene:52050 ; RRID:Addgene_52050) ^75^ and was recloned into a pEG-vector to enable its recombinant expression in HEK293 suspension cells using the BacMam-system. The starting gene contained a K503R point mutation and therefore, all UBR5-constructs used in this study, also contain this point mutation even though being referred to as WT throughout this study. Baculovirus of the respective construct was prepared and used to infect HEK293S cells that were grown to a cellular density of ∼3 Mio cells/mL in Dulbecco’s Modified Eagle Medium (DMEM), supplemented with 10 % fetal calf serum (FCS). In order to help with the expression of Zn-binding domains of UBR5, cells were additionally supplemented with 100 μM ZnSO_4_. Eight hours post-infection, Na-Butyrate was added to a final concentration of 10 mM and cells were grown for 60 h at 30°C. Cells were harvested by centrifugation for 15 min, 450 x g at 4°C, resuspended in lysis buffer and lysed as described above. TwinStrep-tagged GFP-fusion protein was isolated using Strep-affinity chromatography, followed by overnight cleavage with His-tagged PreScission protease. The next day, size exclusion chromatography was carried out. For UBR5^C2765A^ and UBR5^Dimer^, which were used for collection of the apo-UBR5 datasets, the final size exclusion buffer consisted of 25 mM HEPES pH 7.5, 150 mM NaCl, 1 mM DTT. Other UBR5-variants including UBR5^Dimer^, basis for datasets on the distinct ubiquitylation transition states, were purified in a final buffer containing 25 mM HEPES pH 7.5, 150 mM NaCl, 1 mM TCEP. Despite intense efforts to circumvent this, SDS-PAGE gel analysis revealed degradation products of UBR5 after size exclusion chromatography. The identity of these truncated species was confirmed to be UBR5 by mass spectrometry analysis, and remain somewhat catalytically active. The SDS-lane corresponding to the truncations is marked with “*” in all biochemical assays.

### Mass photometry

To determine the oligomeric state of UBR5 and obtain an idea of the molecular mass, mass photometry measurements were performed on the Refeyn OneMP. Mass calibration was achieved by measurement of a protein mixture, providing a range of molecular masses: Ovalbumin (43 kDa), Conalbumin (75 kDa), Aldolase (158 kDa), Ferritin (440 kDa), and Thyroglobulin (669 kDa) in a final concentration of ∼20 nM of each component. Measurements of either UBR5 or UBR5^Dimer^ were carried out by diluting UBR5 to a final concentration of 75 nM in the same buffer used for focus-finding (25 mM HEPES pH 7.5, 150 mM NaCl, 1 mM TCEP). Movies were collected for one minute. Data was then analyzed using the DiscoverMP software and the collected mass calibration as reference.

### Generation of TS1- and TS2-probes

#### Semi-synthesis of Ub-BmDPA

Ub-BmDPA is the building block of our activity-based probes, which aim to visualize the transfer of Ub^D^ from E2 to E3 and subsequently from E3 onto an acceptor Ub. With these approaches, we are trying to mimic native geometry as closely as possible. For this, His-Ub(1-75)-intein-chitin-binding-domain(CBD) was expressed in *E. coli* and cells were lysed in lysis buffer containing 20 mM Tris-HCl pH 6.8, 50 mM NaOAc, 100 mM NaCl, and 2.5 mM PMSF. His-Ub(1-75)-intein-CBD was purified using Ni-NTA affinity chromatography and intein-based cleavage was induced by addition of 100 mM MESNa as previously described ^76^. The resulting His-Ub(1-75)-MESNa was then purified using size exclusion chromatography in a final buffer of 25 mM HEPES pH 6.8., 100 mM NaCl, 20 mM NaOAc. The hydrolysis ratio of His-Ub(1-75)-MESNa was analyzed using liquid chromatography coupled to mass spectrometry (LC-MS) and was taken into account for all further steps. To convert the obtained His-Ub(1-75)-MESNa into chemical proxys, the thioester group modified. His-Ub(1-75)-MESNa (10 mg/mL) was first coupled to 0.4 M (E)-3-[2-(bromomethyl)-1,3-dioxolan-2-yl]prop-2-en-1-amine (BmDPA) (ChiroBlock) in the presence of 1 mM N-hydrosuccinimide in 10 % (v/v) DMSO, and 50 mM HEPES pH 6.8 . After incubating the sample overnight at 30°C, 300 rpm, it was desalted into 25 mM HEPES pH 6.8, 100 mM NaCl and completion of the reaction was confirmed using LC-MS. Next, the product was deprotected by incubating it at a concentration of ∼1 mg/mL in 40 mM p-Toluenesulfonic acid and 54 % (v/v) TFA/H_2_O for 1 h at room temperature. Removal of TFA was achieved by washing the suspension several times with ice-cold diethyl ether. After air-drying the obtained Ub flakes, they were resuspended in 100 mM Na_2_HPO_4_ pH 6.0, 500 mM NaCl, 8 M urea and protein was refolded via dialysis in 20 mM Na_2_HPO_4_ pH 6.0 and 100 mM NaCl overnight at 4°C.

#### Sortase-mediated transpeptidation of Carboxyfluoresceine-PEG5-LPETGG to UBE2D2

Due to its large size of > 300 kDa, it was not possible to rely on the mass shift on an SDS-PAGE gel of UBR5 upon reaction with the UBE2D2-BmDPA-Ub probe. Therefore, we aimed to fluorescently label the E2 enzyme and enable visualization of E3-E2 probe conjugates via fluorescence detection on a Typhoon Scanner. To achieve this, we designed a fluoresceine-labeled LPETGG peptide with a PEG5 linker between fluorophore and sortase recognition sequence (CF-PEG5-LPETGG, MPIB core facility). We then took advantage of the proteolytic TEV-cleavage Gly-Ser remnant at the N-terminus of UBE2D2 during purification. This was recognized by sortase A as a substrate for the transpeptidation reaction with the labeled peptide. Thus, we incubated 6x fold molar excess of labeled-peptide (300 µM) with 50 µM UBE2D2 (C21A/C107A/C111S) and 5 µM His-tagged Sortase A for 1 h at room temperature in 50 mM Tris-HCl pH 8.0, 150 mM NaCl, 10 mM CaCl_2_. Sortase A was removed via His-affinity chromatography and the E2 enzyme-containing flow-through collected. Subsequently, the labeled E2 enzyme was purified via size exclusion chromatography in 50 mM HEPES pH 7.5, 150 mM NaCl.

#### Formation of fluoresceine-labeled and unlabeled UBE2D2-BmDPA-Ub as TS 1 probe

The transition state 1 probe creates a stable mimic of ubiquitin transfer from the catalytic cysteine of UBE2D2 to UBR5’s catalytic cysteine. For the generation of unlabeled UBE2D2-Ub-probe, which was used for cryo-EM, we incubated 100 µM of His-Ub-BmDPA with 5x fold molar excess of UBE2D2^C21A,C107A,C111S^ for 2 h at 30°C. The E2 enzyme was incubated with 1 mM TCEP for 30 min at RT to ensure complete reduction of the catalytic cysteine and desalted into reducing reagent-free buffer before conversion with Ub-BmDPA. Excess E2 enzyme was removed via His-affinity chromatography and further purified by size exclusion chromatography in 25 mM HEPES pH 7.5, 150 mM NaCl. In order to obtain fluorescently-labeled UBE2D2-Ub-probe, utilized to test whether the probe-reactivity is dependent on UBR5’s catalytic cysteine, the molar excess was reversed due to the limited yield of the labeled E2. Consequently, 50 µM of fluoresceine-UBE2D2 was incubated with 5x fold molar excess of Ub-BmDPA (250 µM) for 2 h at 30°C. His-affinity chromatography was skipped and the conjugated probe was directly purified via size exclusion chromatography in 50 mM HEPES pH 7.5, 150 mM NaCl. A synthesis scheme is shown in Extended Data Figure 4 and was prepared using ChemDraw.

#### Generation of Ub^D^-BmDPA-Ub^A^ as TS 2 probe

Ub-BmDPA was also the building block for transition state 2 probe, which creates a stable mimic of transfer of Ub^D^ from UBR5’s catalytic cysteine to Ub^A^’s K48. To generate this reactive probe, the targeted lysine on Ub^A^ (K48) was mutated to Cys. It was essential for Ub^K48C^ to be incubated with fresh reducing reagent (1 mM TCEP) before the conjugation reaction. After desalting into 25 mM HEPES pH 7.5, 150 mM NaCl, Ub^K48C^ was incubated with 5x fold excess of Ub-BmDPA for 1 h at 30°C in the aforementioned buffer. Excess protein was removed using size exclusion chromatography with the final buffer consisting of 25 mM HEPES pH 7.5, 150 mM NaCl. The synthesis scheme is shown in Extended Data Figure 7. Chemical structures were drafted using ChemDraw.

### Generation of probe to mimic E3∼Ub^D^ intermediate state

#### General Procedures for Chemical Synthesis

General reagents were obtained from Sigma Aldrich, Acros and Fluka used as received. Solvents were purchased from Aldrich or BIOSOLVE. Peptide synthesis reagents were purchased from Novabiochem. LC-MS measurements were performed on a Waters Acquity H-class UPLC with a LCT^TM^ ESI-Mass Spectrometer. Samples were run using 2 mobile phases: A = 1 % CH_3_CN, 0.1 % formic acid in water and B = 1 % water and 0.1 % formic acid in CH_3_CN. Data processing was performed using Waters MassLynx Mass Spectrometry Software 4.1 (deconvolution with Maxent1 function).

##### Solid Phase Peptide Synthesis

SPPS was performed on a Syro II MultiSyntech Automated Peptide synthesizer using standard 9-fluorenylmethoxycarbonyl (Fmoc) based solid phase peptide chemistry at 25 μmol scale, using 4 x fold excess of amino acids relative to pre-loaded Fmoc amino acid trityl resin (0.2 mmol/g, Rapp Polymere GmbH).

##### RP-HPLC purifications

Waters preparative RP-HPLC system, equipped with a Waters C18-Xbridge 5 µm OBD (10 x 150 mm) column at a flowrate of 37.5 mL/min. using 3 mobile phases: A: MQ, B: CH_3_CN and C: 1 % TFA in MQ. Prep-HPLC program: Gradient: 0 – 5 min: 5 % B, 5 % C; 5 – 7 min: 5 -> 20% B, 5% C; 7 – 18 min: 20 -> 45 % B, 5 % C. On a Waters C18-Xbridge 5 µm OBD (30 x 150 mm) column at a flowrate of 37.5 mL/min. Pure fractions were pooled and lyophilized.

##### Gel filtration

Size exclusion chromatography was performed on a Sephadex S75 10/300 column (GE Healthcare), using a 20 mM Tris-HCl, 150 mM NaCl buffer, pH 7.6. Appropriate fractions were pooled and concentrated using an Amicon spinfilter (MWCO 10 kDa) to a final concentration of 5.0 mg/mL.

##### (HR)-LC-MS-measurements

High resolution liquid chromatography mass analysis was performed on a Waters Acquity H-Class UPLC system equipped with a Waters ACQUITY Quaternary Solvent Manager(QSM) and Waters ACQUITY FTN AutoSampler. Separation was achieved on a Waters Acquity UPLC Protein BEH C4 column, 300Å, 1,7 uM (2.1 x 50 mm); flow rate = 0.6 mL/min, runtime = 4.55 min, column T = 60°C using 2 mobile phases: A = 0,1% formic acid in water and B = 0,1% formic acid in CH3CN. The products were analyzed by intact MS analysis (MS1) and masses were detected in a range from 550-2000 Da from 2.51 – 4.50 min and were recorded on a Waters XEVO-G2 XS Q-Tof mass spectrometer equipped with an electrospray ion source in positive mode (Capillary Voltage: 0.5 kV, desolvation gas flow: 900 L/h, desolvation gas temperature: 500°C, source temperature: 130 °C, probe angle: 9.5) with a resolution of *R* = 22,000. Data processing was performed using Waters MassLynx Mass Spectrometry Software 4.1 (deconvolution with MaxEnt1 function).

#### Chemical synthesis of Ub-VME and Rhodamine-Ub-VME

Ub(1-75) was synthesized on a Syro II MultiSyntech Automated Peptide synthesizer using standard 9-fluorenylmethoxycarbonyl (Fmoc) based solid phase peptide chemistry on a 25 μmol ^77, 78^ scale. The resin was used as such to prepare Ub-VME or functionalised on the N-terminus by coupling rhodamine. The unmodified or rhodamine-modified Ub was removed from the resin by using 1,1,1,3,3,3-hexafluoropropan-2-ol (HFIP) as described. Gly-VME (5 equiv) was coupled to the C-terminus of Ub using PyBOP (5 equiv), triethylamine (Et_3_N) (10 equiv) in DCM (5 mL) and stirred for 5 h at ambient temperature. Excess Gly-VME was removed by washing the DCM solution with 1 M KHSO_4_. The organic layer was dried with Na_2_SO_4_ and concentrated to dryness in vacuo. To remove the side-chain protecting groups, the residue was taken up in trifluoroacetic acid/triisopropylsilane/water (5 mL; 95:2.5:2.5) and stirred for 3 h at ambient temperature. The reaction mixture was added to a falcon tube containing ice-cold pentane/diethyl ether (1:3; 40 mL), upon which the product precipitated. The precipitate was isolated by centrifugation (1500 x g, 6 min, 4°C) and washed by three cycles of resuspension in ice-cold diethyl ether and centrifugation. Finally, the pellet was taken up in water/acetonitrile/acetic acid (65:25:10), frozen, lyophilized and purified giving Ub-VME as white powder or RhoUbVME as a pink powder. The purity of the peptides was determined by LC-MS analysis and the crude product was purified using RP-HPLC followed by lyophilization of the appropriate fractions.

##### Ub-VME

Deconvoluted ESI MS^+^ (amu) calculated: 8589.6[M+H]^+^; found 8590.0 [M+H]^+^. Rt 1.34 min; HR-MS analysis for C_380_H_631_N_105_O_118_S: [M+ 7H]^7+^ calculated: 1228.24, found: 1227.68, [M+ 8H]^8+^ calculated: 1074.84, found: 1074.35, [M+ 9H]^9+^ calculated: 955.52, found: 954.98, [M+ 10H]^10+^ calculated: 860.07, found: 859.68, [M+ 11H]^11+^ calculated: 781.98, found: 781.62, [M+ 12H]^12+^ calculated: 716.90, found: 716.57.

##### Rho-Ub-VME

Deconvoluted ESI MS^+^ (amu) calculated: 8945.7 [M+H]^+^; found 8943.0 [M+H]^+^. Rt 1.44 min; HR-MS analysis for C_401_H_643_N_107_O_122_S: [M+ 5H]^5+^ calculated: 1790.35, found: 1789.58, [M+ 6H]^6+^ calculated: 1492.13, found: 1491.49, [M+ 7H]^7+^ calculated: 1279.11, found: 1278.56, [M+ 8H]^8+^ calculated: 1119.35, found: 1118.87, [M+ 9H]^9+^ calculated: 995.09, found: 994.66, [M+ 10H]^10+^ calculated: 895.68, found: 895.30.

#### Cryo-EM sample preparation, data collection and processing

##### Tetrameric UBR5^C2768A^

Catalytically inactive UBR5^C2768A^ was freshly purified as described earlier and supplemented with 3-[(3-Cholamidopropyl)-dimethylammonio]-2-hydroxy-1-propansulfonat (CHAPSO) shortly before plunging (8 mM). 3.5 μL sample concentrated to 2.5 mg/mL was applied onto freshly glow discharged R1.2/1.3, Cu 200 mesh holey carbon grids (Quantifoil) using a Vitrobot Mark IV (Thermo Fisher) at 100 % humidity, 4°C. Grids were blotted for 3 seconds, blot force 4 and plunge-frozen into liquid ethane. Data was collected on a Titan Krios transmission electron microscope (TEM) operating at 300 kV, equipped with a post-GIF Gatan K3 Summit direct electron detector in counting mode. Frames were recorded at a nominal magnification of 64,000 x fold and a pixel size of 1.384 Å/pixel at the specimen level with a target defocus range of -2.4 and -0.8 μm and total exposure of 56.28 e^-^/ Å^2^. 10,091 micrographs were collected and alignment and dose-weighing was performed using MotionCorr2 ^79^ followed by CTF estimation using GCTFv1.06 ^80^. Screening datasets of UBR5^C2768A^, collected on a Glacios TEM at 200 kV and equipped with a Gatan K2 detector in counting mode, yielded an initial model. This model was obtained using RELION 3.1.1 and was subsequently used as reference for the Krios dataset. Severe preferred orientation of particles was observed in screening datasets but could be reduced by addition of detergent. 1.1 million particles were picked using template-based picking with Gautomatch (K. Zhang, MRC Laboratory of Molecular Biology, Cambridge). 2D classification, followed by extensive 3D classification, 3D refinements, CTF refinement, particle polishing, and post-processing were performed using RELION 3.1.1. Binning of the particles was lifted stepwise during 3D classifications. Comparison of 3D classes revealed breathing motions of the upper and the lower dimeric unit with respect to each other.

A final map was generated by applying 2 x fold symmetry (C2) during 3D refinement followed by map sharpening using either DeepEMhancer (shown in Figure 1) or post-processing, resulting in a map of 3.7 Å. The intrinsic flexibility of the two dimeric units severely impacted the map quality of the lower half of the map. The processing scheme is depicted in Extended Data Figure 1.

##### _UBR5_^Dimer^

The mutant UBR5^L710D^, herein referred to as UBR5^Dimer^, was supplemented with n-Octyl-β-D-Glucopyranoside (β-OG) at 0.1 % (w/v) shortly before plunging it at a concentration of 1.3 mg/mL. Plunging and data collection were carried out as described for UBR5^C2768A^. Frames were collected at 105,000 x fold magnification with a pixel size of 0.8512 Å/pixel, a target defocus range of -3.0 to -0.5 and a total exposure of 67.8 e^-^/Å^2^. 21,270 micrographs were collected and subjected to alignment and dose-weighing as described above.

Once again, Glacios microscope-derived screening datasets of UBR5^Dimer^, yielded an initial model, that was used as 3D reference and template for particle-picking. Template-based particle picking resulted in 762,722 particles. Data processing was performed using RELION 4.0 ^81^. 2D classification, followed by 3D classification, 3D refinements, and two iterative rounds of CTF refinement as well as particle polishing resulted in a 2.7 Å map after applying C2-symmetry during 3D refinement and map-sharpening using post-processing or DeepEMhancer.

Converting UBR5 to a dimeric species significantly reduced preferred orientations. Polished and refined particles of the final 3D refinement were transferred from RELION to CryoSparc v4.2.0 in order to improve density around regions corresponding to the RLD, and DSD domain. Non-uniform refinement and local refinement was carried out, covering the RLD, a part of the scaffold, and the proximal HECT domain-region ^82, 83^. With the local refinement map in hand, DeepEMhancer implemented in CryoSparc was used to calculate a map with a final resolution of 2.98 Å. While the overall resolution is lower, local resolutions are much higher in the RLD-region compared to the full map. The processing scheme is depicted in Extended Data Figure 3.

##### UBR5^Dimer^∼Ub^D^∼UBE2D2 – TS 1

A fluorescent version of the TS1-probe was used to determine whether probe-reactivity was dependent on the catalytic cysteine of UBR5. For this reason, either UBR5 or the catalytically inactive UBR5^C2768A^ was mixed with ∼5 x fold molar excess of probe and incubated at room temperature for the indicated timepoints. In-gel fluorescence was measured after performing SDS-PAGE to show progression of the probe-reaction (Extended Data Figure 4c). Notably, UBR5^C2768A^ exhibited base-levels of fluorescent signal, possibly due to incomplete proteolytic cleavage of the GFP-tag.

Cryo-EM sample preparation was performed by incubating UBR5^Dimer^ with equimolar amount of K63-linked tetra-ubiquitin chain and 2 x fold molar excess of UBE2D2∼BmDPA∼Ub for 2 h on ice. The reaction mix was plunged without further purification at a concentration of 2 mg/ml. (Note: Reaction of UBR5 with increased excess of E2 probe, longer incubation times, higher temperature and/or purification of the reaction mix did not yield a more homogenous sample as several screening datasets were collected and analysed.) Incubation with n-Octyl-β-D-glucopyranoside (Sigma Aldrich) at a final concentration of 0.1 % (w/v) resulted in a higher density of particles. The detergent was added shortly before applying 3 μL of UBR5^Dimer^∼Ub^D^∼UBE2D2 to freshly glow-discharged holey carbon grids (Quantifoil, R1.2/1.3, 200 mesh). The sample was consequently plunge-frozen into liquid ethane at 95 % humidity and 4°C (blot force 3, blot time 4 s). Data were collected on thin ice with an Arctica electron microscope, equipped with a Falcon III electron detector in linear mode. Movies were collected at a nominal magnification of 73,000x, equaling 1.997 Å/pixel at the specimen level. The target defocus ranged between -3.5 and -1.0 μm and a total exposure of ∼70 e/Å^2^ was distributed over 40 frames.

RELION 4.0. was used for motion correction and dose weighing of 1,740 micrographs. The contrast transfer function was estimated using CTFFIND-4.1 ^84^. The structure of UBR5^Dimer^ with the HECT domains in L-configuration was used as template for picking with Gautomatch and as 3D reference. 1.4 Million particles were extracted (2.1x binned) and subjected to extensive 3D classification. While structures of the initial 3D classification resembled the reference structure with both HECT domains in L-conformation, further classifications displayed either one of the HECTs in an inverted T-conformation with ubiquitin conjugated to the C-lobe. At the same time, sample heterogeneity became apparent as the HECT domains of both UBR5 protomers would adopt either L-, inverted T- or a mix of both conformations. Extensive 3D classification was performed to visualize combinations of HECT conformations and classify out single, stable conformations. This was followed by focused classification with the newly obtained HECT in inverted T-conformation as reference to search for the most stable inverted T-conformation with robust ubiquitin density.

After deriving a clean set of 46,615 particles with one of the two HECT domains fixed in the inverted T-conformation, particles were extracted to full pixel size. Finally, the particle set was refined to 7.3 Å and the 3D reconstruction sharpened using either RELION post-processing or DeepEMhancer. The processing scheme is depicted in Extended Data Figure 5.

##### UBR5^Dimer^∼UbVME – intermediate state

To test whether reactivity of Ub-VME depends on the catalytic cysteine of UBR5, Rho-Ub-VME was incubated with either UBR5 or the catalytically inactive UBR5^C2768A^ at ∼5 x fold molar excess and incubated at room temperature for indicated timepoints. SDS-PAGE with a subsequent in-gel fluorescence measurement exhibited reacted species (Extended Data Figure 4g). Background-signal of UBR5^C2768A^ even in the absence of probe indicates remnants of uncleaved GFP-labeled UBR5.

To prepare the sample for cryo-EM, UBR5^Dimer^ was incubated with equimolar amount of K63-linked tetra-Ubiquitin chain and 10 x fold molar excess of UbVME for 2 h at room temperature. The reaction mix was consequently subjected to size exclusion chromatography (25 mM HEPES pH 7.5, 150 mM NaCl, 0.5 mM TCEP) and peak fractions were concentrated to 1.5 mg/ml.

Shortly before plunging, CHAPSO (Sigma-Aldrich) was added to the protein sample at a final concentration of 8 mM (1x CMC). Subsequently, holey carbon grids (Quantifoil, R1.2/1.3, 200 mesh) were glow discharged, and 3 μL of UBR5^Dimer^∼UbVME were applied to the grid at 95 % humidity and 4°C in a Vitrobot Mark IV (Thermo) and plunge-frozen into liquid ethane (blot force 3, blot time 4 s). Data were collected on medium thick ice with a Glacios electron microscope, equipped with a K2 Summit direct electron detector in counting mode. Movies were collected at a nominal magnification of 22,000x, equaling 1.885 Å/pixel at the specimen level. The target defocus ranged between -2.6 and -0.8 μm and a total exposure of ∼60 e^-^/Å^2^ was distributed over 40 frames.

RELION 4.0 was used for motion correction and dose weighing of 1,808 micrographs. The contrast transfer function was estimated using CTFFIND-4.1, and particles were picked template-free with Gautomatch. 834,722 particles were extracted (2.3x binned) and subjected to 3D classification with the structure of UBR5^Dimer^ serving as 3D reference. After the first round of classification 3D structures displayed robust density for ubiquitin, conjugated to the C-lobe of each protomer. No un-conjugated C-lobe could be observed during processing of the dataset, suggesting complete reaction of UBR5 with UbVME. However, while secondary structures were visible for the HECT domain, ubiquitin density was less defined and of lower resolution-implying flexibility. To investigate this, re-extracted particles to full pixel size of the refined UBR5^Dimer^∼UbVME structure were imported to CryoSparc to perform 3D-Variability Analyses. Default parameters were used for the 3D-VA with structures being low-pass filtered to 9 Å. Substantial movements of the ubiquitin-conjugated C-lobe and the dSBB domain could be observed, justifying lower resolution of those domains.

The final masked refinement of UBR5^Dimer^∼UbVME with 197,281 particles yielded a 3D reconstruction at 5.3 Å and was sharpened using either RELION post-processing or DeepEMhancer. The processing scheme is depicted in Extended Data Figure 6.

##### UBR5∼Ub^D^∼Ub^A^ TS 2

Transfer of Ub^D^ from the catalytic cysteine of UBR5 to Ub^A^ was mimicked by conjugation of UBR5^Dimer^ with a ∼50 x fold molar excess of the Ub^D^-BmDPA-Ub^A^ in 25 mM HEPES pH 7.5, 150 mM NaCl, 1 mM TCEP for 2 h at RT. UBR5-Ub^D^-Ub^A^ was purified using size exclusion chromatography in a final buffer of 25 mM HEPES pH 7.5, 150 mM NaCl, 1 mM TCEP. The sample was pooled and concentrated to 0.6 mg/mL and subsequently supplemented with CHAPSO (final detergent concentration 8 mM). Plunging was performed as described for UBR5^C2768A^ and a dataset was collected on a Glacios screening microscope with a target defocus of -3.0 and -0.3 μm and a total exposure of ∼60 e^-^/Å^2^ partitioned into 40 frames. RELION 3.1.1 was used for motion correction and dose weighing of 705 micrographs. The contrast transfer function was estimated using GCTF. A low-pass filtered map of UBR5^Dimer^ was used as template for picking with Gautomatch and as initial 3D reference. ∼300,000 particles were picked and subjected to 2D and 3D classification. 3D refinement with subsequent sharpening using post-processing or DeepEMhancer resulted in a 8.3 Å map that had significant additional density next to the C-lobe of both HECT domains. Its overall architecture resembled UBR5^Dimer^∼Ub^D^, however, additional density next to the C-lobe and Ub^D^ could accomodate Ub^A^ and the UBA domain.

To improve map quality of the HECT domain, conjugated with Ub^A^, Ub^D^, including the Ub^A^-neighboring UBA domain, a second dataset was collected on UBR5∼Ub^D^∼Ub^A^ on a Titan Krios electron microscope with a magnification of 105,000 x fold, a pixel size of 0.8512 Å/pixel, and a defocus range of -2.2 to -0.6. 17,689 micrographs were collected and alignment as well as dose-weighing were performed as described for other datasets. Template-based picking with the UBR5^Dimer^ model resulted in 1.7 million particles. 2D classification as well as several rounds of 3D classification with and without image alignment were performed using RELION 3.1.1. A mask was created covering the HECT domain, Ub^A^ and Ub^D^, as well as the neighboring UBA domain and two rounds of masked 3D classifications were carried out. Next, classified particles and the 3D reconstruction were imported to CryoSparc. There, non-uniform-refinement followed by local refinement, focusing on the HECT domain, Ub^A^, Ub^D^, and the UBA domain calculated a resolution of 3.3 Å. The map was sharpened using DeepEMhancer.

#### Model building and refinement

##### Model for tetrameric UBR5

An initial model of UBR5 was generated using the AlphaFold2 server. The obtained model was split into smaller parts in order to be docked into the density map obtained for UBR5^C2768A^ using UCSF Chimera. This allowed for determination of residue L710 for mutagenesis to disrupt the interaction connecting both “U”-shaped units.

##### Structure for UBR5^Dimer^

The high-resolution map of UBR5^Dimer^ allowed precise building of the protein backbone including side-chains in most parts of the structure using Coot. Due to the low resolution for the dSBB domain, the barrels could not be built but were docked into the structure using the AlphaFold2-model instead. Unfortunately, we were not able to determine how residues 1523-1773 connect at the heterodimerization interface without unambiguity. For this reason, we assigned them as four separate chains in the coordinates. However, we note that AlphaFold2 predicts the structural connection between these, and protomers were therefore assigned on this basis for figures. To build the RLD and DSD domain, the focused map of UBR5^Dimer^ was used due to better map quality in those regions. Since density for the connection of the DSD domain to the scaffold was missing, it could originate from either monomer and was therefore kept as separate chain. However, the closer distance of the monomer in trans suggests that the DSD domain originates there and integrates into the other monomer to increase the dimerization-interface.

In early refinement cycles, two-fold symmetry was applied to the structure of a monomer to obtain the structure of a dimer. Lastly, several rounds of real-space refinement were performed using PHENIX. The atomic model of UBR5^Dimer^ was validated using Molprobity^85^. A complete summary of data collection and refinements statistics were provided in Extended Data Table 1.

##### Model for UBR5^Dimer^∼ Ub^D^∼ UBE2D2– TS 1

To generate a model for TS1 (UBR5^Dimer^∼ Ub^D^∼ UBE2D2), the structure of UBR5^Dimer^ was split into several parts: the scaffold that could be readily docked into the density, the HD domain and the N-lobe of the HECT domain, which had to be tilted slightly compared to the apo-UBR5-structure. The HECT domain C-lobe had to be massively rearranged. UBE2D2∼Ub was extracted from a preexisting-structure where it was bound to NEDD4L (PDB: 3JVZ). Using UCSF Chimera, these components were fitted into the obtained map. Note that UBE2D2 in 3JVZ has the catalytic cysteine mutated to serine to facilitate trapping by oxyester-formation. Instead, UBE2D2 used in our study contains the catalytic cysteine, however, the three remaining cysteines are mutated as follows: C21A, C107A, C111S ^86^. Since the obtained density for TS1 only had one HECT domain positioned in the inverted T-conformation (“HECT 1”) and the other HECT domain presumably being a mix of L- and inverted T-conformation (“HECT 2”), the model only applies to HECT domain 1.

##### Model of UBR5^Dimer^∼Ub^D^ – intermediate state

An initial model for the intermediate state was made based on a previously published structure of Rsp5∼Ub x Sna3 (PDB: 4LCD) and the structure of UBR5^Dimer^. The PDB was fitted into the density for HECT domain 2 using UCSF Chimera. Subsequently, Ub^D^ was extracted from this PDB and docked into the density individually. The final model for the intermediate state, shown in Figure 3, was built based on the structure for TS2. Since the HECT∼Ub^D^-conformation of the TS2 structure fitted nicely into the density, these parts were docked into the density together with the remaining parts of the UBR5^Dimer^ structure. Since the EM-map does not exhibit density for the C-terminus of UBR5 and C-terminus of Ub^D^ (also differing in the reactive group compared to TS2), these parts were truncated for the final model of UBR5^Dimer^∼Ub^D^. Even though the map reveals clear density for Ub^D^ on both HECT domains, our model only focuses on one HECT domain bound to Ub^D^ (“HECT 2”).

##### Structure of UBR5∼Ub^D^∼Ub^A^ – transition state 2

To build the structure mimicking transition state 2, the model of UBR5∼Ub^D^, as well as the crystal structure showing UBR5’s UBA domain bound to ubiquitin (PDB: 2QHO) were docked into the focused map. The map quality largely allowed building of the HECT domain including both ubiquitins, Ub^A^ and Ub^D^, on a side-chain level using Coot. In this model, a K48C-mutation of Ub^A^ was introduced. There is clear density for residues 2796-2798 (Phe-Gly-Phe), although the precise locations of side-chains were ambiguous. For Phe2796, density was smeared, suggesting potential conformations for this residue. For Phe2798, lack of clear visibility for the subsequent, terminal residue precludes absolute determination if the density corresponds to the main chain or the side-chain. The density is tentatively assigned to the side-chain, based on the biochemical effect of mutations, but its unambiguous placement will require future studies. Side-chains of Arg72 and Arg74 of Ub^D^ were slightly moved to allow placement of UBR5’s C-terminus. Placement of the Arg54 rotamer in Ub^A^ was carried out in consideration biochemical phenotypes caused by mutagenesis of E2287 and R54. However, future studies will be required to determine the precise location of these side-chains. Even though, the bonds of the applied probe were not visible in the density, we know the geometry of the chemical probe. The three-way cross-link was built between UBR5 Cys 2768, G75 of Ub^D^ and C48 of Ub^A^ as shown in Figure 4A. The final model was subjected to several iterations of real-space refinement in PHENIX. Validation of the atomic model was performed using Molprobity.

#### Generation of di-Ubs and higher Ub chains

##### Enzymatic assembly of K6, K11, K48 and K63-linked di-ubiquitin species

K6, K11, K48, and K63-linked di-ubiquitins were prepared using enzymatic assembly. All of them were generated by employing distinct E2 or E3 enzymes, with tagless Ub purified as described previously.

Formation of K6 linked di-Ub was achieved by incubating 2.5 mM Ub with 0.1 μM E1, 0.6 μM UBE2L3, 10 μM NleL in the presence of 10 mM ATP in 40 mM Tris-HCl pH 8.8, 10 mM MgCl_2_, 1 mM DTT for 3 h at 37°C. The reaction was quenched with 10 mM DTT and K48 linked ubiquitin chains that were generated as a byproduct, were removed by subsequent incubation with 2 μM OTUB1 for 3 h at 37°C.

K11 linked Ub chains were obtained by incubating 0.5 mM Ub with 0.25 μM E1, and 5 μM Ube2S-UBA-IsoT in the presence of 10 mM ATP for 2 h at 37°C.

To generate K48 linked di-Ub, 2.5 mM Ub were incubated with 1 μM E1, and 25 μM UBE2R1 in the presence of 10 mM ATP for 3 h at 37°C. The reaction was quenched by adding 10 mM DTT and 1 μM AMSH.

K63 linked di-Ub was generated by incubating 1 mM Ub with 0.5 μM E1, 8 μM Ube2N and 8 μM Ube2V1 in 10 mM ATP, 40 mM Tris-HCl pH 8.5, 10 mM MgCl_2_, 0.5 mM DTT at 37°C for 30 min before stopping the reaction by addition of 10 mM DTT (final concentration).

Different chain lengths of the various chain types were then separated using iterative rounds of ion exchange chromatography followed by size exclusion chromatography in a final buffer of 25 mM HEPES pH 7.5.

##### Chemical synthesis of K27, K29 and K33-linked di-ubiquitin species

K27, K29, and K33linked di-Ub was prepared using analogues methods as described in ^78^ in short:

The γ-thiolysine-Ub_p_ (StBu protected) was dissolved in DMSO (15 mg/100 µL) and added to 8 M Gdn-HCl/100 mM phosphate buffer, pH 7.6 (final concentration 7.5 mg/mL) supplemented with 100 mM TCEP and reacted at 37°C. After LC-MS analysis revealed complete deprotection of the thiolysine 1 equivalent of Ub^D^-thioester dissolved in DMSO (15 mg/100 µL) and 8 M Gdn.HCl/100 mM phosphate buffer, pH 7.6 (final concentration 7.5 mg/mL) and 50 mM MPAA were added. The pH was readjusted to 7.6 and the reaction was reacted for 16 h at 37°C. RP-HPLC purification was followed by lyophilization of the appropriate fractions.

Desulfurization was achieved by dissolving in DMSO (concentration 15 mg/100 µL) and subsequent dilution into 8 M Gdn. HCl/100 mM phosphate buffer at pH 7.6 (final concentration 5 mg/mL) and 150 mM TCEP. VA044 (25 mg/mL) and GSH (25 mg/mL) were added again, the pH was adjusted to 7.0. The mixture was allowed to react for 16 hours at 37°C followed by RP-HPLC purification and lyophilization of the appropriate fractions.

The lyophilized di-Ub was dissolved in DMSO (concentration 7.5 mg/100 µL) and diluted into 20 mM Tris-HCl pH 7.6, 150 mM NaCl (7.5 mg/mL) and purified on an S75 16/600 Sephadex size exclusion column. Appropriate fractions were collected and pooled followed by spin-filtration (Amicon 3 kDa MWCO) to concentrate the sample to 5.0 mg/mL. The aliquots were snap frozen and stored at -80°C until further use.

Deconvoluted ESI MS^+^ (amu) calcd: 17093.7, found 17094.4, rt: 1.36 min. HR-MS analysis for C_757_H_1258_N_210_O_235_S : [M+ 10H]^10+^ calculated: 1710.3376, found: 1710.3276, [M+ 11H]^11+^ calculated: 1554.9436, found: 1554.9438, [M+ 12H]^12+^ calculated: 1425.4490, found: 1425.4441, [M+ 13H]^13+^ calculated: 1315.8766, found: 1315.8804, [M+ 14H]^14+^ calculated: 1221.9574, found: 1221.9567, [M+ 15H]^15+^ calculated: 1140.5608, found: 1140.5573, [M+ 16H]^16+^ calculated: 1069.3387, found: 1069.3364, [M+ 17H]^17+^ calculated: 1006.4957, found: 1006.4905, [M+ 18H]^18+^ calculated: 950.6353, found: 950.6315, [M+ 19H]^19+^ calculated: 900.6549, found: 900.6536, [M+ 20H]^20+^ calculated: 855.6725, found: 855.6719, [M+ 21H]^21+^ calculated: 814.9742, found: 814.9700, [M+ 22H]^22+^ calculated: 777.9758, found: 777.9737, [M+ 23H]^23+^ calculated: 744.1946, found: 744.1901, [M+ 24H]^24+^ calculated: 713.2284, found: 713.2257, [M+ 25H]^25+^ calculated: 684.7396, found: 684.7383.

### Biochemical assays

#### Pulse-chase format: Di-ubiquitin synthesis assay

Di-ubiquitin synthesis assays in a pulse-chase format were carried out to examine the catalytic effects of different UBR5-versions and distinct Ub^A^-mutants. The assays were performed with fluorescently labeled donor Ub and unlabeled acceptor Ub. The employed E2 was UBE2D2 as it showed significantly higher reactivity towards UBR5 than UBE2L3^87^. 20 μM of UBE2D2 was incubated with 30 μM fluorescent donor Ub^K48R^ in the presence of 0.5 μM UBA1 in a buffer containing 25 mM HEPES pH 7.5, 150 mM NaCl, 5 mM MgCl_2_, 2 mM ATP, and 0.04 mg/mL BSA for 30 min at room temperature. Loading of the E2 was quenched with 50 mM EDTA.

During the chase-reaction, UBR5 variants were mixed with distinct unlabeled acceptor ubiquitin mutants to test how well these can collaborate. In general, the thioester-linked E2∼*Ub^D^ was diluted into a mix of E3 (0.2 μM final) and the respective Ub^A^-6xHis (2 μM final) in 25 mM HEPES pH 7.5, 150 mM NaCl, 1 mM DTT (“chase buffer”) to a final concentration of 0.2 μM unless stated otherwise. Samples were taken after the indicated times and the reaction was quenched by adding non-reducing SDS-PAGE buffer.

##### UBR5 variants

Different properties of UBR5 were tested with tetrameric UBR5-versions that don’t contain the L710D mutation. Tetrameric UBR5 was used to test the effect of mutating the catalytic cysteine (Figure 1b), the C-lobe–Ub^D^-interaction site A2790 (Figure 2e), SDA^mut^ (Extended Data Figure 4e), different variations of the C-terminal UBR5-tail (Figure 5c), the UBA-mutation L224D (Figure 5d), and the LOL-mutant (Figure 5g). To test the following properties, dimeric UBR5 (containing the L710D mutation) was utilized: how well acceptor mutants containing D58-mutations (Figure 5c), or mutations in the UBA-Ub^A^-interface (Figure 5d) can get modified by UBR5, as well as what effect a UBR5-mutation in the C-lobe-Ub^A^ (Figure 5e), or the N-lobe-Ub^A^-interface (Figure 5f) has.

Samples were taken at the indicated time points and mixed with SDS-PAGE loading buffer. After performing SDS-PAGE, in-gel fluorescence was scanned using a Typhoon Imager (Amersham). Intensity of the scans was increased and the gels were cropped subsequently.

##### UBR5 size truncations

UBR5 truncations with various lengths were also tested for their ability to form di-Ubs (Figure 1i). The constructs used during the chase-reaction comprised either the entire C-terminal HECT domain, including the inserted MLLE domain (“HECT”) or with the MLLE domain being replaced by a 6 amino acid long linker (“HECT^ΔMLLE^”), the UBA domain fused N-terminally by a 15 amino acid long linker to the complete HECT domain with the MLLE domain inserted (“UBA-HECT”), UBR5^Dimer^ or UBR5 respectively. To test these constructs, the respective E3-variant (0.5 μM final concentration) was mixed with Ub^A^-6xHis (5 μM final concentration) and 1 μM E2∼*Ub^D^. Samples were taken after 0.3, 2, and 10 min and mixed with SDS-PAGE loading buffer. In-gel fluorescence was scanned subsequent to SDS-PAGE. The intensity was increased during figure preparation.

##### Determination of linkage-specifity

To test linkage-specificity of UBR5 and UBR5^Dimer^, the chase-reaction was prepared by mixing 0.2 μM of UBR5 or UBR5^Dimer^ with 2 μM of different acceptor ubiquitins in the chase buffer. The acceptor ubiquitin contained either all lysines (“WT”), no lysines (all lysines were mutated to arginines = “K0”), or only one distinct lysine (“6”,”11”,”27”,”29”,”33”,”48”, or”63”), and all other lysines were mutated to arginines. For the chase-reaction, 0.2 μM E2∼*Ub^D^ was added to the prepared mix and samples were taken after 1 min. The reaction was stopped by adding SDS-PAGE loading buffer supplemented with 100 mM DTT. Subsequent to performing SDS-PAGE, in-gel fluorescence was scanned. The gels were cropped during figure preparation and the intensity was manually increased (Extended Data Figure 2l).

##### E2-UBR5 N-lobe interface mutants

To test whether mutating the E2 in the E2-N-lobe interface would affect the di-Ub-formation, the point mutation F62A of UBE2D3 and UBE2D3 WT were used during the pulse-reaction. 20 μM of the respective E2 were incubated with 30 μM fluorescent donor Ub^K48R^ and 0.5 μM UBA1 in a buffer containing 25 mM HEPES pH 7.5, 150 mM NaCl, 5 mM MgCl_2_, 2 mM ATP, and 0.04 mg/mL BSA for 30 min at room temperature. Loading of the E2 was quenched with 50 mM EDTA. Subsequently, 0.2 μM E2∼*Ub^D^ were added to a mix of 0.2 μM UBR5 and 2 μM Ub^A^-6xHis. Samples were taken at the indicated times and the reaction was stopped by mixing with SDS-PAGE loading buffer. In-gel fluorescence was scanned after performing SDS-PAGE. During figure preparation, intensity of the gels was increased and they were cropped.

##### pH influence on LOL-mutant

Whether a varying pH influences the transpeptidation-activity of UBR5 was also addressed employing the di-Ub synthesis assay. The pulse-reaction was performed as previously described to not influence E2-loading. Instead of performing the chase-reaction in 25 mM HEPES pH 7.5, 150 mM NaCl, 1 mM DTT, different buffer components were now used. For the reactions performed at pH 6.8, 7.5, 8.0, and 8.8, 25 mM Tris-HCl adjusted to the respective pH, was used. The remaining components – 150 mM NaCl, 1 mM DTT – were maintained in all reactions, regardless of the pH. To facilitate an even higher pH, the buffer compound had to be changed: pH 9.5 was achieved using CAPSO and pH 11 was realized using CAPS. 0.2 μM UBR5 or UBR5^D2283–2287A^ was mixed with 2 μM Ub^A^-6xHis in the respective buffer. E2∼*Ub^D^ was diluted into this sample by a ratio of 1:20 to not affect the pH significantly. Samples were taken by adding SDS-PAGE loading buffer without supplemented DTT after 60 sec and visualization of the result was performed using SDS-PAGE followed by fluorescent imaging. The intensity of the scan was increased and gels were cropped for clarity.

##### Pulse-chase format: Tri-Ub synthesis assay

To test how well UBR5 can modify differently linked di-Ubs, and to see whether this correlates with the accessibility of the respective lysine on Ub^A^, a tri-Ub synthesis assay was performed in Figure 4J. The pulse-reaction was performed as described for the di-Ub synthesis assay with fluorescently-labeled Ub^K48R^ being loaded onto UBE2D2. The reaction was quenched by addition of 50 mM EDTA. A pulse-mix was generated by mixing UBR5^WT^ (0.2 μM final concentration) and di-Ub linked via the respective lysine (2 μM final concentration) in 25 mM HEPES pH 7.5, 150 mM NaCl, 1 mM DTT. As described earlier, M1-linked di-Ubs were obtained by linear fusions, K6, K11, K48, or K63-linked di-Ubs were generated enzymatically, and K27, K29, or K33-linked di-Ubs were generated by chemical synthesis. Subsequently, E2∼*Ub^D^ was added to the pulse-mix at a final concentration of 0.2 μM. Samples were taken after 20 sec and mixed with SDS-PAGE loading buffer supplemented with a final concentration of 100 mM DTT. After performing SDS-PAGE, the gel was scanned for in-gel fluorescence. The image was cropped during figure-preparation and the intensity was adjusted.

#### Pulse-chase format: Autoubiquitylation/free Ub chain formation assay

Whether both, UBR5 and UBR5^Dimer^ can form polyubiquitin chains to a comparable extent, a pulse-chase assay was performed with fluoresceine-labeled WT Ub that could serve as both, donor, and acceptor. The pulse-reaction was performed by incubating 30 μM labeled Ub with 20 μM UBE2D2, and 0.5 μM E1 in 2 mM ATP, 10 mM MgCl2, 25 mM HEPES pH 7.5, 150 mM NaCl, and 0.04 mg/mL BSA for 30 min at room temperature. The reaction was stopped by addition of 50 mM EDTA and the chase-reaction was performed by mixing 0.2 μM of the respective UBR5-version with 1 μM E2∼*Ub. Samples were taken at the indicated time points and mixed with reducing SDS-loading buffer. Using fluorescent scanning subsequently to SDS-PAGE, the result was visualized and the intensity of the scan was increased during the process of figure-making.

##### Multi-turnover format: Polyubiquitylation

To test the polyubiquitylation-activity of the SDA-mutant UBR5^H1362–1364D^, a multi-turnover assay was performed. For this, fluoresceine-labeled WT Ub was mixed at 20 μM with 5 μM UBE2D2, and 0.5 μM of UBR5 or SDA^mut^ in 2 mM ATP, 5 mM MgCl_2_, 25 mM HEPES pH 7.5, 150 mM NaCl, and 0.04 mg/mL BSA. Addition of 0.5 μM E1 started the reaction and samples were taken at the indicated time points, mixed with reducing SDS-loading buffer. The result was visualized using fluorescent scanning subsequently to SDS-PAGE.

## REFERENCES

1 Komander, D. & Rape, M. The ubiquitin code. Annu Rev Biochem 81, 203–229, doi:10.1146/annurev-biochem-060310-170328 (2012).

2 Husnjak, K. & Dikic, I. Ubiquitin-binding proteins: decoders of ubiquitin-mediated cellular functions. Annu Rev Biochem 81, 291–322, doi:10.1146/annurev-biochem-051810-094654 (2012).

3 Chau, V. et al. A multiubiquitin chain is confined to specific lysine in a targeted short-lived protein. Science 243, 1576–1583, doi:10.1126/science.2538923 (1989).

4 Spence, J., Sadis, S., Haas, A. L. & Finley, D. A ubiquitin mutant with specific defects in DNA repair and multiubiquitination. Mol Cell Biol 15, 1265–1273, doi:10.1128/MCB.15.3.1265 (1995).

5 Galan, J. M. & Haguenauer-Tsapis, R. Ubiquitin lys63 is involved in ubiquitination of a yeast plasma membrane protein. EMBO J 16, 5847–5854, doi:10.1093/emboj/16.19.5847 (1997).

6 Kirisako, T. et al. A ubiquitin ligase complex assembles linear polyubiquitin chains. EMBO J 25, 4877–4887, doi:10.1038/sj.emboj.7601360 (2006).

7 Rahighi, S. et al. Specific recognition of linear ubiquitin chains by NEMO is important for NF-kappaB activation. Cell 136, 1098–1109, doi:10.1016/j.cell.2009.03.007 (2009).

8 Meyer, H. J. & Rape, M. Enhanced protein degradation by branched ubiquitin chains. Cell 157, 910–921, doi:10.1016/j.cell.2014.03.037 (2014).

9 Yau, R. G. et al. Assembly and Function of Heterotypic Ubiquitin Chains in Cell-Cycle and Protein Quality Control. Cell 171, 918–933 e920, doi:10.1016/j.cell.2017.09.040 (2017).

10 Liu, C., Liu, W., Ye, Y. & Li, W. Ufd2p synthesizes branched ubiquitin chains to promote the degradation of substrates modified with atypical chains. Nat Commun 8, 14274, doi:10.1038/ncomms14274 (2017).

11 Ohtake, F., Tsuchiya, H., Saeki, Y. & Tanaka, K. K63 ubiquitylation triggers proteasomal degradation by seeding branched ubiquitin chains. Proc Natl Acad Sci U S A 115, E1401–E1408, doi:10.1073/pnas.1716673115 (2018).

12 French, M. E., Koehler, C. F. & Hunter, T. Emerging functions of branched ubiquitin chains. Cell Discov 7, 6, doi:10.1038/s41421-020-00237-y (2021).

13 Kolla, S., Ye, M., Mark, K. G. & Rape, M. Assembly and function of branched ubiquitin chains. Trends Biochem Sci 47, 759–771, doi:10.1016/j.tibs.2022.04.003 (2022).

14 Berndsen, C. E. & Wolberger, C. New insights into ubiquitin E3 ligase mechanism. Nat Struct Mol Biol 21, 301–307, doi:10.1038/nsmb.2780 (2014).

15 Buetow, L. & Huang, D. T. Structural insights into the catalysis and regulation of E3 ubiquitin ligases. Nat Rev Mol Cell Biol 17, 626–642, doi:10.1038/nrm.2016.91 (2016).

16 Zheng, N. & Shabek, N. Ubiquitin Ligases: Structure, Function, and Regulation. Annu Rev Biochem 86, 129–157, doi:10.1146/annurev-biochem-060815-014922 (2017).

17 Christensen, D. E., Brzovic, P. S. & Klevit, R. E. E2-BRCA1 RING interactions dictate synthesis of mono- or specific polyubiquitin chain linkages. Nat Struct Mol Biol 14, 941–948, doi:10.1038/nsmb1295 (2007).

18 Kim, H. C. & Huibregtse, J. M. Polyubiquitination by HECT E3s and the determinants of chain type specificity. Mol Cell Biol 29, 3307–3318, doi:10.1128/MCB.00240-09 (2009).

19 Rotin, D. & Kumar, S. Physiological functions of the HECT family of ubiquitin ligases. Nat Rev Mol Cell Biol 10, 398–409, doi:10.1038/nrm2690 (2009).

20 Mattiroli, F. & Sixma, T. K. Lysine-targeting specificity in ubiquitin and ubiquitin-like modification pathways. Nat Struct Mol Biol 21, 308–316, doi:10.1038/nsmb.2792 (2014).

21 Scheffner, M., Nuber, U. & Huibregtse, J. M. Protein ubiquitination involving an E1-E2-E3 enzyme ubiquitin thioester cascade. Nature 373, 81–83, doi:10.1038/373081a0 (1995).

22 Huibregtse, J. M., Scheffner, M., Beaudenon, S. & Howley, P. M. A family of proteins structurally and functionally related to the E6-AP ubiquitin-protein ligase. Proc Natl Acad Sci U S A 92, 2563–2567, doi:10.1073/pnas.92.7.2563 (1995).

23 Wenzel, D. M., Lissounov, A., Brzovic, P. S. & Klevit, R. E. UBCH7 reactivity profile reveals parkin and HHARI to be RING/HECT hybrids. Nature 474, 105–108, doi:10.1038/nature09966 (2011).

24 Stieglitz, B. et al. Structural basis for ligase-specific conjugation of linear ubiquitin chains by HOIP. Nature 503, 422–426, doi:10.1038/nature12638 (2013).

25 Branigan, E., Plechanovova, A., Jaffray, E. G., Naismith, J. H. & Hay, R. T. Structural basis for the RING-catalyzed synthesis of K63-linked ubiquitin chains. Nat Struct Mol Biol 22, 597–602, doi:10.1038/nsmb.3052 (2015).

26 Pan, M. et al. Structural insights into Ubr1-mediated N-degron polyubiquitination. Nature 600, 334–338, doi:10.1038/s41586-021-04097-8 (2021).

27 Cotton, T. R. et al. Structural basis of K63-ubiquitin chain formation by the Gordon-Holmes syndrome RBR E3 ubiquitin ligase RNF216. Mol Cell 82, 598–615 e598, doi:10.1016/j.molcel.2021.12.005 (2022).

28 Nakasone, M. A. et al. Structure of UBE2K-Ub/E3/polyUb reveals mechanisms of K48-linked Ub chain extension. Nat Chem Biol 18, 422–431, doi:10.1038/s41589-021-00952-x (2022).

29 Scheffner, M. & Kumar, S. Mammalian HECT ubiquitin-protein ligases: biological and pathophysiological aspects. Biochim Biophys Acta 1843, 61–74, doi:10.1016/j.bbamcr.2013.03.024 (2014).

30 Kee, Y. & Huibregtse, J. M. Regulation of catalytic activities of HECT ubiquitin ligases. Biochem Biophys Res Commun 354, 329–333, doi:10.1016/j.bbrc.2007.01.025 (2007).

31 Lorenz, S. Structural mechanisms of HECT-type ubiquitin ligases. Biol Chem 399, 127–145, doi:10.1515/hsz-2017-0184 (2018).

32 Kamadurai, H. B. et al. Insights into ubiquitin transfer cascades from a structure of a UbcH5B approximately ubiquitin-HECT(NEDD4L) complex. Mol Cell 36, 1095–1102, doi:10.1016/j.molcel.2009.11.010 (2009).

33 Kamadurai, H. B. et al. Mechanism of ubiquitin ligation and lysine prioritization by a HECT E3. Elife 2, e00828, doi:10.7554/eLife.00828 (2013).

34 Maspero, E. et al. Structure of a ubiquitin-loaded HECT ligase reveals the molecular basis for catalytic priming. Nat Struct Mol Biol 20, 696–701, doi:10.1038/nsmb.2566 (2013).

35 Nair, R. M. et al. Reconstitution and Structural Analysis of a HECT Ligase-Ubiquitin Complex via an Activity-Based Probe. ACS Chem. Biol. 16, 1615–1621, doi:10.1021/acschembio.1c00433 (2021).

36 Fajner, V., Maspero, E. & Polo, S. Targeting HECT-type E3 ligases – insights from catalysis, regulation and inhibitors. FEBS Lett 591, 2636–2647, doi:10.1002/1873-3468.12775 (2017).

37 Shearer, R. F., Iconomou, M., Watts, C. K. & Saunders, D. N. Functional Roles of the E3 Ubiquitin Ligase UBR5 in Cancer. Mol Cancer Res 13, 1523–1532, doi:10.1158/1541-7786.MCR-15-0383 (2015).

38 Liao, L. et al. E3 Ubiquitin Ligase UBR5 Drives the Growth and Metastasis of Triple-Negative Breast Cancer. Cancer Res 77, 2090–2101, doi:10.1158/0008-5472.CAN-16-2409 (2017).

39 Qiao, X. et al. UBR5 Is Coamplified with MYC in Breast Tumors and Encodes an Ubiquitin Ligase That Limits MYC-Dependent Apoptosis. Cancer Res 80, 1414–1427, doi:10.1158/0008-5472.CAN-19-1647 (2020).

40 Schukur, L. et al. Identification of the HECT E3 ligase UBR5 as a regulator of MYC degradation using a CRISPR/Cas9 screen. Sci Rep 10, 20044, doi:10.1038/s41598-020-76960-z (2020).

41 Oh, E. et al. Gene expression and cell identity controlled by anaphase-promoting complex. Nature 579, 136–140, doi:10.1038/s41586-020-2034-1 (2020).

42 Song, M. et al. Tumor derived UBR5 promotes ovarian cancer growth and metastasis through inducing immunosuppressive macrophages. Nat Commun 11, 6298, doi:10.1038/s41467-020-20140-0 (2020).

43 Xiang, G. et al. UBR5 targets tumor suppressor CDC73 proteolytically to promote aggressive breast cancer. Cell Death Dis 13, 451, doi:10.1038/s41419-022-04914-6 (2022).

44 Kaisari, S. et al. Role of ubiquitin-protein ligase UBR5 in the disassembly of mitotic checkpoint complexes. Proc Natl Acad Sci U S A 119, doi:10.1073/pnas.2121478119 (2022).

45. Mark, K. G., et al. Orphan quality control shapes network dynamics and gene expression. bioRxiv, 2022.2011.2006.515368, doi:10.1101/2022.11.06.515368 (2022).

46 Kozlov, G. et al. Structural basis of ubiquitin recognition by the ubiquitin-associated (UBA) domain of the ubiquitin ligase EDD. J Biol Chem 282, 35787–35795, doi:10.1074/jbc.M705655200 (2007).

47 Kozlov, G., Menade, M., Rosenauer, A., Nguyen, L. & Gehring, K. Molecular determinants of PAM2 recognition by the MLLE domain of poly(A)-binding protein. J Mol Biol 397, 397–407, doi:10.1016/j.jmb.2010.01.032 (2010).

48 Matta-Camacho, E., Kozlov, G., Menade, M. & Gehring, K. Structure of the HECT C-lobe of the UBR5 E3 ubiquitin ligase. Acta Crystallogr Sect F Struct Biol Cryst Commun 68, 1158–1163, doi:10.1107/S1744309112036937 (2012).

49 Wang, F. et al. Structure of the human UBR5 E3 ubiquitin ligase. bioRxiv, 2022.2010.2031.514604, doi:10.1101/2022.10.31.514604 (2022).

50. Hodáková, Z., et al. Cryo-EM structure of the chain-elongating E3 ligase UBR5. bioRxiv, 2022.2011.2003.515015, doi:10.1101/2022.11.03.515015 (2022).

51 Tasaki, T. et al. The substrate recognition domains of the N-end rule pathway. J Biol Chem 284, 1884–1895, doi:10.1074/jbc.M803641200 (2009).

52 Wang, F. et al. Structure of the human UBR5 E3 ubiquitin ligase. Structure 31, 541–552 e544, doi:10.1016/j.str.2023.03.010 (2023).

53 Lim, N. S. et al. Comparative peptide binding studies of the PABC domains from the ubiquitin-protein isopeptide ligase HYD and poly(A)-binding protein. Implications for HYD function. J Biol Chem 281, 14376–14382, doi:10.1074/jbc.M600307200 (2006).

54 Kane, E. I. et al. Redefining the catalytic HECT domain boundaries for the HECT E3 ubiquitin ligase family. Biosci Rep 42, doi:10.1042/BSR20221036 (2022).

55 Bremm, A., Freund, S. M. & Komander, D. Lys11-linked ubiquitin chains adopt compact conformations and are preferentially hydrolyzed by the deubiquitinase Cezanne. Nat Struct Mol Biol 17, 939–947, doi:10.1038/nsmb.1873 (2010).

56 Horn-Ghetko, D. et al. Ubiquitin ligation to F-box protein targets by SCF-RBR E3-E3 super-assembly. Nature 590, 671–676, doi:10.1038/s41586-021-03197-9 (2021).

57 Nuber, U. & Scheffner, M. Identification of determinants in E2 ubiquitin-conjugating enzymes required for hect E3 ubiquitin-protein ligase interaction. J Biol Chem 274, 7576–7582, doi:10.1074/jbc.274.11.7576 (1999).

58 Borodovsky, A. et al. Chemistry-based functional proteomics reveals novel members of the deubiquitinating enzyme family. Chem Biol 9, 1149–1159, doi:10.1016/s1074-5521(02)00248-x (2002).

59 Punjani, A. & Fleet, D. J. 3D variability analysis: Resolving continuous flexibility and discrete heterogeneity from single particle cryo-EM. J Struct Biol 213, 107702, doi:10.1016/j.jsb.2021.107702 (2021).

60 Salvat, C., Wang, G., Dastur, A., Lyon, N. & Huibregtse, J. M. The -4 phenylalanine is required for substrate ubiquitination catalyzed by HECT ubiquitin ligases. J Biol Chem 279, 18935–18943, doi:10.1074/jbc.M312201200 (2004).

61 Burch, T. J. & Haas, A. L. Site-directed mutagenesis of ubiquitin. Differential roles for arginine in the interaction with ubiquitin-activating enzyme. Biochemistry 33, 7300–7308, doi:10.1021/bi00189a035 (1994).

62 Lee, I. & Schindelin, H. Structural insights into E1-catalyzed ubiquitin activation and transfer to conjugating enzymes. Cell 134, 268–278, doi:10.1016/j.cell.2008.05.046 (2008).

63. Mao, J., et al. Structural Visualization of HECT-E3 Ufd4 accepting and transferring Ubiquitin to Form K29/K48-branched Polyubiquitination on N-degron. bioRxiv, 2023.2005.2023.542033, doi:10.1101/2023.05.23.542033 (2023).

64 Jackl, M. et al. beta-Sheet Augmentation Is a Conserved Mechanism of Priming HECT E3 Ligases for Ubiquitin Ligation. J Mol Biol 430, 3218–3233, doi:10.1016/j.jmb.2018.06.044 (2018).

65 Hunkeler, M. et al. Solenoid architecture of HUWE1 contributes to ligase activity and substrate recognition. Mol Cell 81, 3468–3480 e3467, doi:10.1016/j.molcel.2021.06.032 (2021).

66 Grabarczyk, D. B. et al. HUWE1 employs a giant substrate-binding ring to feed and regulate its HECT E3 domain. Nat Chem Biol 17, 1084–1092, doi:10.1038/s41589-021-00831-5 (2021).

67 Chen, Z. et al. A Tunable Brake for HECT Ubiquitin Ligases. Mol Cell 66, 345–357 e346, doi:10.1016/j.molcel.2017.03.020 (2017).

68 Zhu, K. et al. Allosteric auto-inhibition and activation of the Nedd4 family E3 ligase Itch. EMBO Rep 18, 1618–1630, doi:10.15252/embr.201744454 (2017).

69 Wang, Z. et al. A multi-lock inhibitory mechanism for fine-tuning enzyme activities of the HECT family E3 ligases. Nat Commun 10, 3162, doi:10.1038/s41467-019-11224-7 (2019).

70 Attali, I. et al. Ubiquitylation-dependent oligomerization regulates activity of Nedd4 ligases. EMBO J 36, 425–440, doi:10.15252/embj.201694314 (2017).

71 Weber, J., Polo, S. & Maspero, E. HECT E3 Ligases: A Tale With Multiple Facets. Front Physiol 10, 370, doi:10.3389/fphys.2019.00370 (2019).

72 Sherpa, D. et al. GID E3 ligase supramolecular chelate assembly configures multipronged ubiquitin targeting of an oligomeric metabolic enzyme. Mol Cell 81, 2445–2459 e2413, doi:10.1016/j.molcel.2021.03.025 (2021).

73 Mohamed, W. I. et al. The human GID complex engages two independent modules for substrate recruitment. EMBO Rep 22, e52981, doi:10.15252/embr.202152981 (2021).

74 Scott, D. C. et al. Two Distinct Types of E3 Ligases Work in Unison to Regulate Substrate Ubiquitylation. Cell 166, 1198–1214 e1124, doi:10.1016/j.cell.2016.07.027 (2016).

75 Gudjonsson, T. et al. TRIP12 and UBR5 suppress spreading of chromatin ubiquitylation at damaged chromosomes. Cell 150, 697–709, doi:10.1016/j.cell.2012.06.039 (2012).

76 Li, G., Liang, Q., Gong, P., Tencer, A. H. & Zhuang, Z. Activity-based diubiquitin probes for elucidating the linkage specificity of deubiquitinating enzymes. Chem Commun (Camb) 50, 216–218, doi:10.1039/c3cc47382a (2014).

77 de Jong, A. et al. Ubiquitin-based probes prepared by total synthesis to profile the activity of deubiquitinating enzymes. Chembiochem 13, 2251–2258, doi:10.1002/cbic.201200497 (2012).

78 El Oualid, F., et al. Chemical synthesis of ubiquitin, ubiquitin-based probes, and diubiquitin. Angew Chem Int Ed Engl 49, 10149–10153, doi:10.1002/anie.201005995 (2010).

79 Zheng, S. Q. et al. MotionCor2: anisotropic correction of beam-induced motion for improved cryo-electron microscopy. Nat Methods 14, 331–332, doi:10.1038/nmeth.4193 (2017).

80 Zhang, K. Gctf: Real-time CTF determination and correction. J Struct Biol 193, 1–12, doi:10.1016/j.jsb.2015.11.003 (2016).

81 Kimanius, D., Dong, L., Sharov, G., Nakane, T. & Scheres, S. H. W. New tools for automated cryo-EM single-particle analysis in RELION-4.0. Biochem J 478, 4169–4185, doi:10.1042/BCJ20210708 (2021).

82 Punjani, A., Zhang, H. & Fleet, D. J. Non-uniform refinement: adaptive regularization improves single-particle cryo-EM reconstruction. Nat Methods 17, 1214–1221, doi:10.1038/s41592-020-00990-8 (2020).

83 Punjani, A., Rubinstein, J. L., Fleet, D. J. & Brubaker, M. A. cryoSPARC: algorithms for rapid unsupervised cryo-EM structure determination. Nat Methods 14, 290–296, doi:10.1038/nmeth.4169 (2017).

84 Rohou, A. & Grigorieff, N. CTFFIND4: Fast and accurate defocus estimation from electron micrographs. J Struct Biol 192, 216–221, doi:10.1016/j.jsb.2015.08.008 (2015).

85 Williams, C. J. et al. MolProbity: More and better reference data for improved all-atom structure validation. Protein Sci 27, 293–315, doi:10.1002/pro.3330 (2018).

86 Baek, K. et al. NEDD8 nucleates a multivalent cullin-RING-UBE2D ubiquitin ligation assembly. Nature 578, 461–466, doi:10.1038/s41586-020-2000-y (2020).

87 Pao, K. C. et al. Activity-based E3 ligase profiling uncovers an E3 ligase with esterification activity. Nature 556, 381–385, doi:10.1038/s41586-018-0026-1 (2018).

